# Shared Lineage, Distinct Outcomes: Yap and Taz Loss Differentially Impact Schwann and Olfactory Ensheathing Cell Development Without Disrupting GnRH-1 Migration

**DOI:** 10.1101/2025.02.13.638196

**Authors:** Ed Zandro M. Taroc, Enrico Amato, Alexis Semon, Nikki Dolphin, Briane Beck, Sophie Belin, Yannick Poitelon, Paolo E. Forni

## Abstract

Olfactory Ensheathing Cells (OECs) are glial cells originating from the neural crest, critical for bundling olfactory axons to the brain. Their development is crucial for the migration of Gonadotropin-Releasing Hormone-1 (GnRH-1) neurons, which are essential for puberty and fertility. OECs have garnered interest as potential therapeutic targets for central nervous system lesions, although their development is not fully understood.

Our single-cell RNA sequencing of mouse embryonic nasal tissues suggests that OECs and Schwann cells share a common origin from Schwann cell precursors yet exhibit significant genetic differences. The transcription factors Yap and Taz have previously been shown to play a crucial role in Schwann cell development. We used *Sox10*-Cre mice to conditionally ablate *Yap* and *Taz* in migrating the neural crest and its derivatives. Our analyses showed reduced Sox10+ glial cells along nerves in the nasal region, altered gene expression of SCs, melanocytes, and OECs, and a significant reduction in olfactory sensory neurons and vascularization in the vomeronasal organ. However, despite these changes, GnRH-1 neuronal migration remained unaffected.

Our findings highlight the importance of the Hippo pathway in OEC development and how changes in cranial neural crest derivatives indirectly impact the development of olfactory epithelia.

## Introduction

The nasal area of mammals comprises cartilages, bones, mesenchyme, and glia of neural crest origin and various sensory and endocrine neurons of placodal origin. Olfactory Ensheathing Cells (OECs), also known as the olfactory Schwann cells or nasal glia, interact with the neurons of the olfactory system (Chuah & Au, 1991). The OECs are crucial in guiding olfactory neuron development and facilitating axonal bundling (Pingault et al., 2013; Zhou, Li, Wu, Yu, & Xia, 2017). For several decades, it was believed that the olfactory placode, which gives rise to pioneer olfactory neurons, terminal nerve neurons, olfactory neurons, vomeronasal neurons, sustentacular cells, Gonadotropin-Releasing Hormone 1 (GnRH-1) neurons (Amato, Taroc, & Forni, 2024; Barraud et al., 2010; Wray, Grant, & Gainer, 1989) - could give rise to OECs cells (Couly & Le Douarin, 1985). However, studies in chickens, fish, and mice indicate that the glia of the olfactory system derives from Neural Crest Cells (NCCs) (Aguillon et al., 2018; Barraud et al., 2010; Forni, Taylor-Burds, Melvin, Williams, & Wray, 2011; Saxena, Peng, & Bronner, 2013). NCCs are multipotent precursor cells that form at the dorsal-most portion of the closing neural tube during embryonic development. NCCs subsequently migrate throughout the embryo, differentiating into diverse cell types, including bones and cartilages, melanocytes, sensory neurons, autonomic and enteric neurons, and all subtypes of peripheral glia known as Schwann cells (SCs) (Erickson, Kameneva, & Adameyko, 2023; Munoz & Trainor, 2015). SCs originate from Schwann cell precursors (SCPs), a transient cell population that shares a multipotent “hub” state with the migratory NCCs (M. E. Kastriti et al., 2022; Stierli & Sommer, 2022). SCs of the peripheral nervous system include myelinating cells as well as a variety of non-myelinating SCs, including the Remak SCs, enteric glia, and terminal SCs, which localize to the neuromuscular junction. OECs share several molecular features with the non-myelinating SCs (Barber & Lindsay, 1982; Doucette, 1984; Franklin & Barnett, 2000). Yet, despite SCs and OECs having overlapping molecular features, the two cell types have been proposed to form distinct progenitors (Perera et al., 2020). In addition, a recent study suggested that the OECs might originate from precursors of the mesenchyme (Perera et al., 2020) and bioinformatic data indicate that the mesenchyme and glial cells may have alternative programs (Erickson et al., 2023), leaving the developmental trajectories of the OECs an open-ended question.

Sox10 is a transcription factor belonging to the Sox family and contains the HMG-box, which is indispensable for the proper development of various cell types derived from the neural crest (Aoki et al., 2003; Britsch et al., 2001; Honore, Aybar, & Mayor, 2003). Disruptions in OEC development after *Sox10* loss-of-function cause olfactory and terminal nerve defects that disrupt the migration of the GnRH-1 neurons from the nasal area to the brain in humans and mice (Barraud, St John, Stolt, Wegner, & Baker, 2013; Pingault et al., 2013; Taroc, Naik, et al., 2020; Zhou et al., 2017); these have been proposed as underlying conditions for Kallmann syndrome. In addition, haploinsufficiency of the *Sox10* gene in humans is associated with type IV Waardenburg syndrome (Pingault et al., 1998). This condition is characterized by several features, including sensorineural hearing loss, hair pigment abnormalities, lateral displacement of the eyes (known as dystopia canthorum), deafness, the absence of OECs, impaired sense of smell (anosmia), and hypogonadotropic hypogonadism. The latter condition results from the defective migration of GnRH-1 neurons (Barraud et al., 2013; Pingault et al., 2013). Despite their pivotal role in the development and function of olfactory and GnRH-1 systems, the cell biology and developmental trajectories of the OECs have been largely overlooked.

YAP (encoded by *Yap1*, shorten to *Yap*) and TAZ (encoded by *Wwtr1*, shorten to *Taz*) are transcriptional co-regulators of the Hippo pathway, which controls multicellular development by integrating chemical and mechanical signals (Cobbaut et al., 2020; Hillmer & Link, 2019; Jafarinia et al., 2024; Manning, Kroeger, & Harvey, 2020; Rogg et al., 2023; Rognoni & Walko, 2019). When the Hippo pathway is inactive, *Yap* and *Taz* can translocate into the nucleus and regulate gene transcription. Previous studies revealed that *Yap* and *Taz* are required for SC development, myelination, and peripheral nerve repair (Deng et al., 2017; Grove et al., 2017; Grove, Lee, Zhao, & Son, 2020; Hong et al., 2024; Jeanette et al., 2021; Y. Poitelon et al., 2016). Moreover, *Wnt1*-Cre-driven conditional manipulation of *Yap* and *Taz* in presumptive pre-mandatory NCCs suggests significant roles for these factors in cranial NCC proliferation, survival, cellular differentiation (J. Wang et al., 2016), and melanocyte gene expression (Manderfield et al., 2015).

In this article, we present our findings using single-cell RNA sequencing of developing mouse noses at E14.5. Our study suggests that during embryonic development, the OECs and SCs in the peripheral nervous system share similar developmental trajectories, as both cell types appear to originate from SCPs in the nasal area. Additionally, we found strong expression of *Yap* and *Taz* in developing OECs. *Sox10*-Cre conditional ablation of *Yap* and *Taz* (*Yap*^cHet^;*Taz*^cKO^) in neural crest cells determined defective gene expression in the nasal mesenchyme, melanocyte, defective SCs, and abnormal OEC development. In *Yap*^cHet^;*Taz*^cKO^ mutants, we found a reduction in olfactory and vomeronasal organ neurons, defective vascularization of the disorganization of the olfactory/vomeronasal axon bundles, but, surprisingly no significant impact on GnRH-1 neuronal migration.

## Results

### Single-cell RNA Sequencing of E14.5 mouse nose identifies developing OECs

To get a full scope of the populations within the developing nose, we dissociated and performed single-cell RNA sequencing of whole mouse noses from embryonic day (E) 14.5 mouse embryos. Unsupervised clustering using the ‘R’ package Seurat allowed us to identify distinct cell types based on the expression of defining markers (Fig. 1A,C). In the developing olfactory epithelia, we categorized *Pax6*-, *Ascl1*-, and *Tubb3*-positive cells as the developing neuronal population (Cau, Gradwohl, Fode, & Guillemot, 1997; Kim, Leung, Reed, & Johnson, 2007; Roskams, Cai, & Ronnett, 1998) (Fig. 1A,B,D), while *Foxa1*-positive cells as non-sensory/respiratory-epithelium (Forni, Bharti, Flannery, Shimogori, & Wray, 2013; Taroc, Katreddi, & Forni, 2020) (Fig. 1A,C,D). *Col1a1*/*Alx1*-positive cells were identified as mesenchyme, cartilage, and or bone, with a subset of cells within the cluster also expressing *Sox10* (Fig. 1A,B,C,D). *Pecam1*-positive cells were labeled as vasculature (Muller, Ratti, McDonnell, & Cohn, 1989). Immune cells were labeled based on *C1qa* (Kishore, Thielens, & Gaboriaud, 2016), *Acta2* for fibroblasts (Hinz, Celetta, Tomasek, Gabbiani, & Chaponnier, 2001), and *Actg2* for muscle cells (Halim et al., 2015). Notably, among the different cell clusters, we identified putative SCPs derivatives/glial cells based on *Fabp7* and *Sox10* enrichment (Barraud et al., 2010; Forni et al., 2011) (Fig. 1A,D). As mentioned earlier, *Sox10* mRNA was expressed in the developing nasal cartilage and sparse non-neuronal cells in the developing olfactory epithelium; these are presumptive developing Bowman gland cells (Perera et al., 2020) (Fig. 1A,B,C,D).

**Figure 1.**
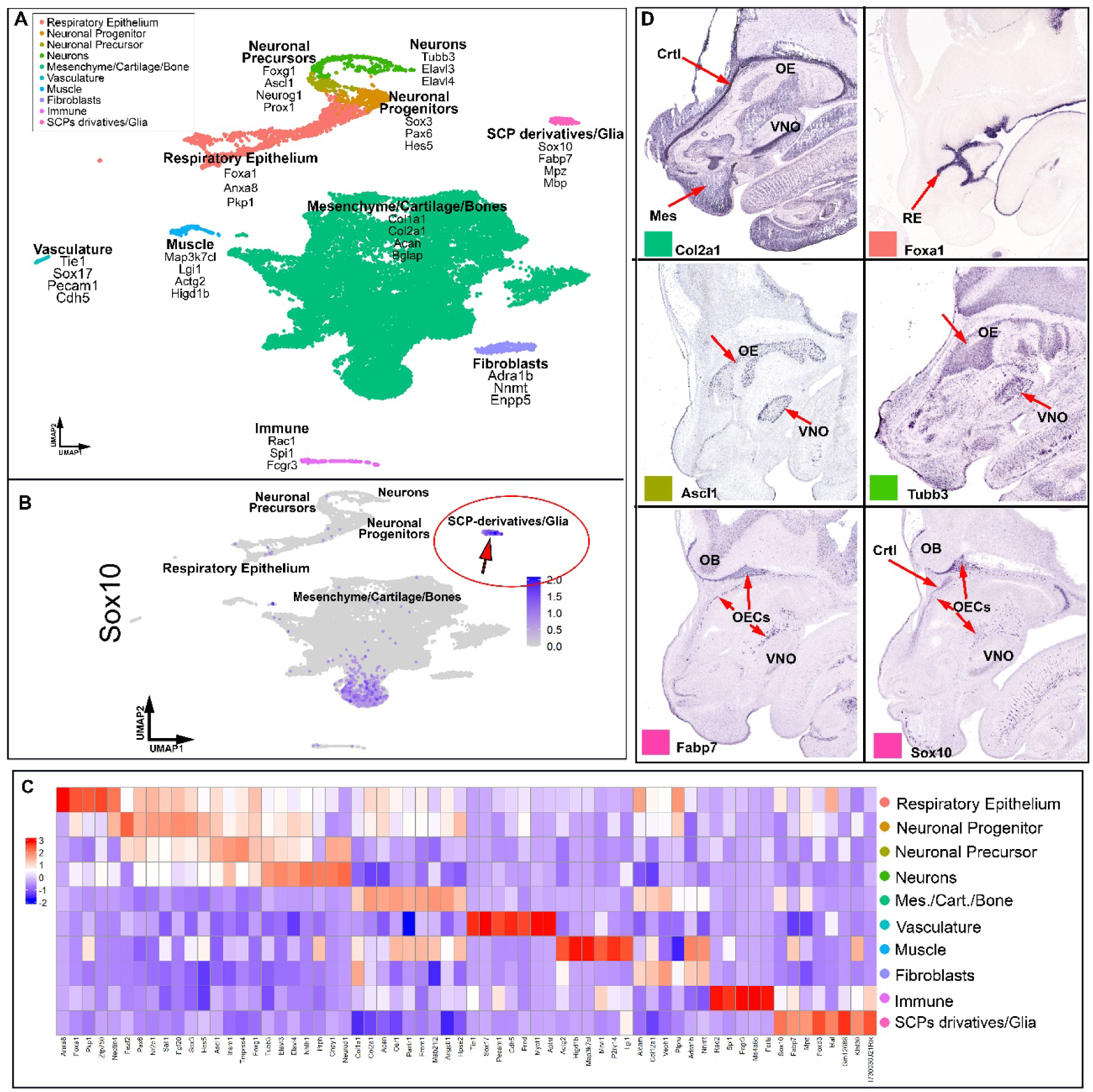
Single-cell sequencing of the noses of E14.5 C57BL/6 embryos. A) UMAP displaying cell clusters. The color code for the presumptive cell types is indicated in the legend. Some of the top genes for each cell type are shown on the UMAP. B) Feature plot for Sox10 expression on the same UMAP as in A. Sox10 is enriched in the cluster of presumptive Schwann Cell Precursor (SPC) derivatives/glia (red arrow).C) Heatmap illustrating differentially expressed genes across clusters. D) In situ hybridization showing the expression of Col2a1 in the nasal mesenchyme (Mes) and cartilage (Crtl); foxa1 in the respiratory epithelium; Ascl1, and Tubb3 in the developing olfactory epithelium (OE) and vomeronasal organ (VNO); Fabp7 and Sox10 in the presumptive olfactory ensheathing cells. (OECs) along the neurons emerging from OE and VNO and surrounding the olfactory bulb (OB). Sox10 is also visible in the nasal cartilage (Crtl). In situ hybridization images were sourced from the GenePaint database (https://gp3.mpg.de).

### Subclustering the Schwann cell precursors and their derivatives suggests a common origin for Schwann cell OECs. (fig2)

Previous studies suggested that the OECs derive from cells of the nasal mesenchyme (Perera et al., 2020). However, the group of *Sox10*-positive cells, comprising the SCPs derivatives/glial cells, formed an isolated cluster not in a continuum with the nasal mesenchyme (Fig.1 A,B). Sub-clustering the *Sox10* enriched SCPs derivatives/glial cell cluster (Fig. 1A, B; 2A,B) determined the formation of 4 subclusters that we classified as 4 cell types: presumptive melanocytes (rich in *Mitf* expression), SCPs (*Ki67*/proliferative), SCs (low *Fabp7*), and OECs (high in *Fabp7*) (Perera et al., 2020).

**Figure 2.**
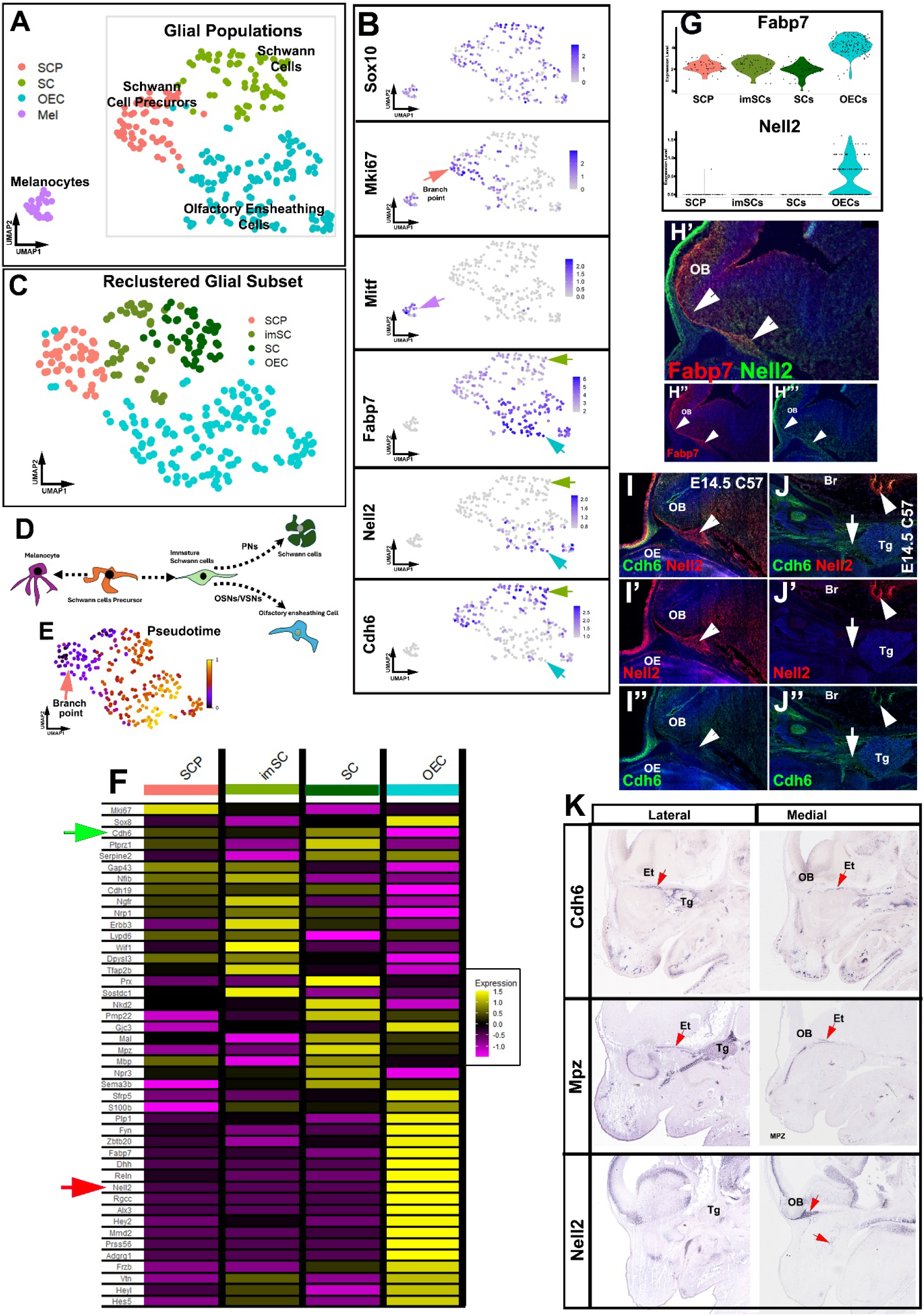
The re-clustering of the Sox10-enriched cluster “Schwann Cell Precursor (SPC) derivatives/glia" indicates that the OECs form from SCPs. **A)** The Sox10-enriched cluster includes four cell types: Schwann Cell Precursors (SCPs), Melanocytes, Schwann Cells (SCs), and Olfactory Ensheathing Cells (OECs). SCs and OECs seem to form a dichotomy beginning with SCPs. B) Feature plots display enriched genes in specific subclusters; the arrow colors indicate the clusters of the same colors in A. C) Re-clustering of (A) without melanocytes implies the presence of an intermediate cell population of immature SCs (imSCs). D) A model proposing that the OECs associated with olfactory/vomeronasal neurons (OSNs/VSNs) and the Schwann cells (SCs) of the peripheral nerves (PNs) both originate from Schwann cell precursors (SCPs). E) RNA Velocity indicates that OECs and SCs differentiate starting from SCPs; darker colors indicate a less-differentiated state, while orange-yellow signifies more differentiated states. F) Heatmap illustrating enriched genes in SCPs, imSCs, SCs, and OECs. G) Violin plots demonstrate Fabp7 and Nell2 enrichment in OECs. H) Nell2 and Fabp7 colocalization (arrowheads) in OECs surrounding the olfactory bulb (OB). I-J”) Cdh6 and Nell2 immunoreactivity appear to be mutually exclusive, with Nell2 enriched in OECs and Cdh6 in SCs along the trigeminal projections (Tg). K) In situ hybridizations on lateral and medial sections of E14.5 mouse heads showing Cdh6, Mpz, and Nell2. Cdh6 and Mpz RNA are visible in the ethmoidal projections (Et) of the trigeminal ganglion (Tg), while Nell2 is not found in the trigeminal projections but in the OECs in the nose and surrounding the olfactory bulb (OB) (images are from gene paint).

To delve deeper into the gene expression patterns of the glial cells, we focused exclusively on isolating and re-clustering the group of cells that gave rise to the dichotomy and excluded the presumptive melanocytes (light gray box in Fig. 2A; Fig. 2C). Utilizing scVelo, we conducted RNA Velocity analysis (Fig. 2E). We implemented a pseudo-temporal reconstruction in Python, allowing us to track the dynamic gene expression along the developmental trajectories of the two branches (Fig. 2E). Examining differential gene expression (heatmap, Fig. 2F) highlighting key markers previously identified in the literature as specific to SCPs, SCs, or OECs (Perera et al., 2020; Rich et al., 2018). We found that the "branch point" expressed the recently discovered SCP marker genes such as *Dlx1/2* and *Itga4* (Maria Eleni Kastriti et al., 2022) (Fig. 2E), as well as the broadly accepted markers for SCPs such as *Gap43*, *Nfib*, and *Cdh19* (K. R. Jessen & Mirsky, 2019; Perera et al., 2020) (Fig. 2F,G). Conversely, the branch with low *Fabp7* expression was found to be positive for gene markers that align with SCs such as *Mpz*, *Mbp*, and *Mal* (Liu et al., 2015; Perera et al., 2020) (Fig. 2C,G), while the branch exhibiting high *Fabp7* (Fig. 2F,G) expression corresponds to OEC marker genes such as *Reln*, *Nell2,* and *Alx3* (Perera et al., 2020; Taroc, Lin, Tulloch, Jaworski, & Forni, 2019). Notably, after refining the clustering to only include the glial populations, the SC branch could be further defined as immature Schwann Cells (imSCs) and mature Schwann Cells (SCs) (Fig 2. C). The imSC population can be genetically defined by a reduction of proliferative markers such as *Mki67* and the mixed expression of SCP markers such as *Gap43*, *Nfib*, *Ngfr*, with mature SC markers like *Gjc3*, and *S100b* (Fig. 2F)(Kristjan R Jessen & Mirsky, 2005; K. R. Jessen & Mirsky, 2019; M. E. Kastriti et al., 2022) indicating that this is a transitional stage during SC development.

To validate the specificity of some of these markers, we conducted immunofluorescent staining that showed us that the OECs along the olfactory nerve layer (notched arrow) are immunoreactive for Fabp7 and Nell2 but not for Cdh6 (Fig. 2 H-I) (Perera et al., 2020; Taroc et al., 2019), while the SCs along the trigeminal ganglion (arrow) are strongly immunoreactive for Cdh6 (Fig. 2H-J’) but not for Nell2. This is further validated by ISH data from Genepaint where *Cdh6* and *Mpz* mRNA can be localized to the trigeminal (Tg) and etomoidal (Et) projections, while *Nell2* mRNA is found surrounding the olfactory bulb and olfactory epithelium/vomeronasal organ (Fig. 2K).

### Hippo-pathway effector molecules *Yap*, *Taz*, and *Tead1/2* are expressed broadly in the developing nasal glial populations. (fig3)

YAP and TAZ regulate gene expression by integrating external and internal signals that control diverse transcriptional programs. YAP and TAZ can shuttle between the cytoplasm and the nucleus, depending on the stimulus. In the nucleus, they pair with DNA-binding factors from the TEAD family to regulate gene expression(Vassilev, Kaneko, Shu, Zhao, & DePamphilis, 2001; Zhang et al., 2008).

Previous studies have shown that the loss of the Hippo effector molecules YAP and TAZ in SCs, led to defects in their differentiation and function (Jeanette et al., 2021; Yannick Poitelon et al., 2016). To investigate whether YAP and TAZ are expressed in the SCPs, SCs, and OECs we looked at our single-cell RNA sequencing data. Feature plots and violin plots for Yap1, Taz/Wwtr1,Tead1 and Tead2 show expression in all clusters within the glial populations, suggesting a role for Hippo signaling in OEC development. (Fig. 3A,B).

**Figure 3.**
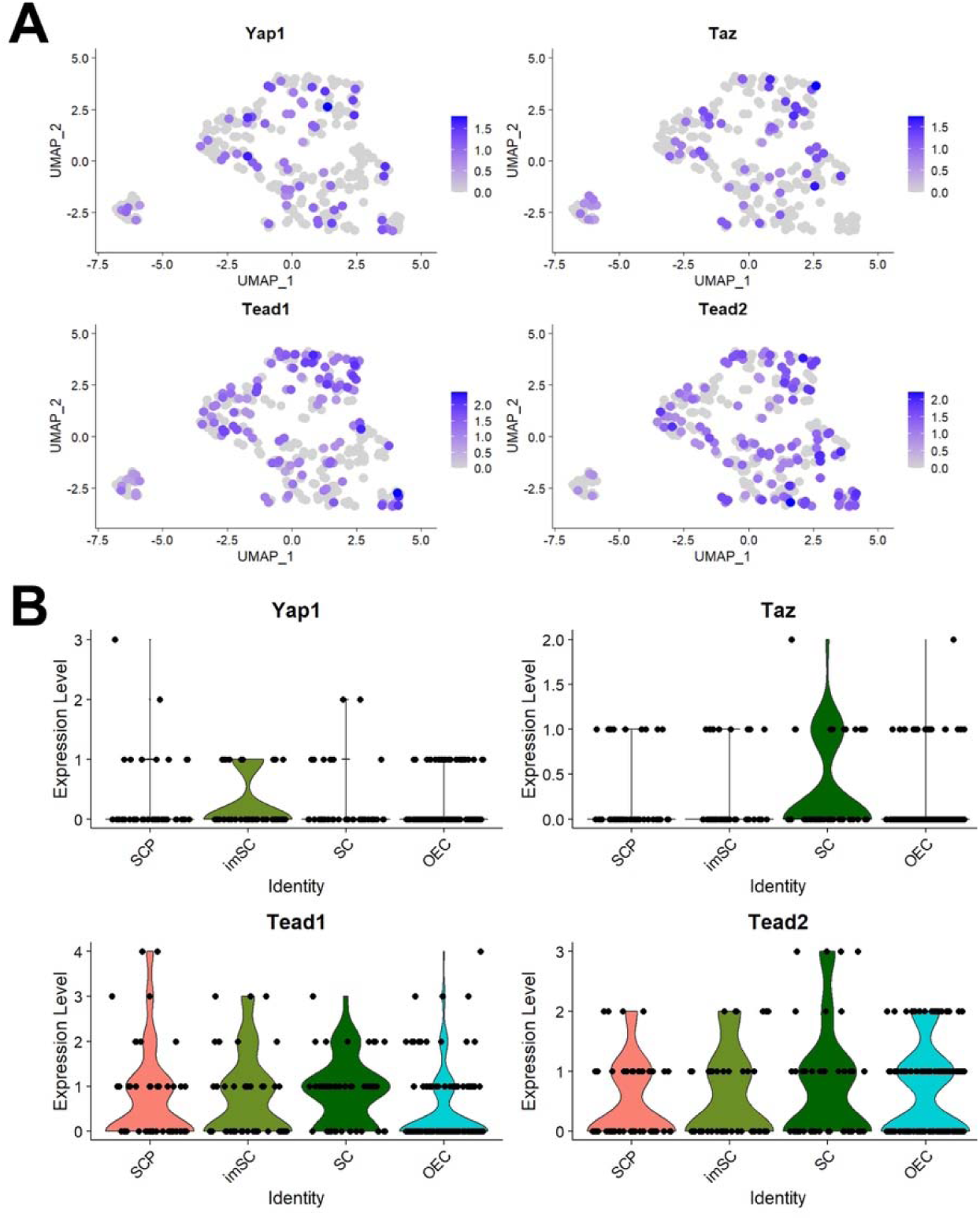
Yap Taz, Tead1/2 are expressed in the developing nasal glia. Feature plots (A) and violin plots (B) show Yap1 and Taz, Tead1 and Tead2 expression in SCPs, imSC, SCs, and OECs.

### *Sox10*-Cre mouse line gives a more restricted NCC recombination than the pre-migratory *Wnt1*-Cre2 mouse line. (fig4)

*Wnt1* is expressed in pre-migratory neural crest as they form in the dorsal neural tube but is not expressed by migratory NCC or its derivatives. Consistent with this, single-cell sequencing of the nose at E14.5 showed no cells expressing *Wnt1* (Fig. 4A). By crossing the widely used *Wnt1*-Cre2 mouse line with the Ai14 Rosa26 reporter, we observed an extensive recombination in the olfactory bulb, OECs, nasal mesenchyme, and in cells within the olfactory/vomeronasal epithelia. In both the olfactory epithelium (OE) and vomeronasal organ (VNO), we detected neuronal cells positive for recombination and for the neuronal markers Elav-like protein 3 antigen C and Elav protein 4 antigen D (HuC/D) as well as cells only positive for recombination. In contrast, *Sox10*, which is expressed later in the migratory NCC and is downregulated in most derivatives, except SCPs, SCs, melanocytes, OECs, and some mesenchymal cells of the frontonasal region (Fig. 1C). Notably, the intermixing of pre-migratory NCCs with cells forming cranial placodes, as well as the integration of presumptive migratory NCC with olfactory epithelia, have been previously proposed and reported (Forni et al., 2011; Forni & Wray, 2015; Katoh et al., 2011; Murdoch, DelConte, & Garcia-Castro, 2010; Saxena et al., 2013; Whitlock, 2004).

**Figure 4.**
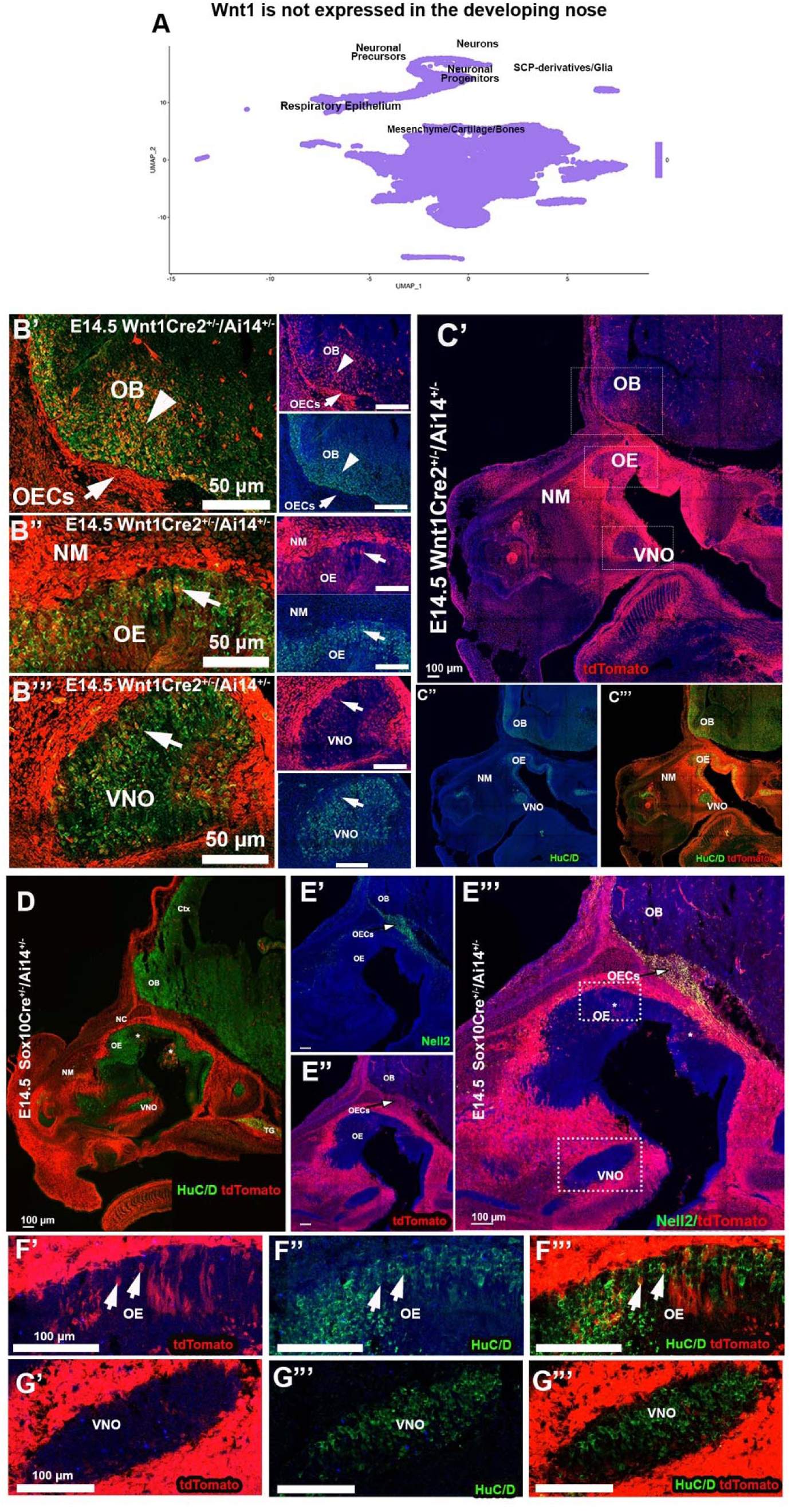
Pre-migratory NC Wnt1Cre2 broadly recombines in cells in non-NC derivatives, while the post-migratory Sox10Cre gives more faithful NC tracing. A) UMAP,single-cell sequencing at E14.5 reveals that Wnt1 is not expressed in the cells of the developing nose. B-B’’’) Immunofluorescence with anti-tdTomato (red), HuC/D (green), and DAPI (blue). B’) Wnt1-Cre2 recombination occurs in the cells of the olfactory bulbs (OB) and olfactory ensheathing cells (OECs). B”) Recombination in neurons and non-neuronal cells of the olfactory epithelium (OE); B”) Recombination in neurons of the vomeronasal organ (VNO). C’-C”) Immunofluorescence with anti-tdTomato (Red), HuC/D (green), and DAPI (blue) displays broad recombination in the OB, OE, VNO, and nasal mesenchyme (NM). D) Sox10Cre/Ai14 mouse at E14.5; immunofluorescence with anti-tdTomato (RED), HuC/D (green), and DAPI (blue) indicates recombination in the nasal mesenchyme (NM) and in sparse cells of the OE. E’-E”’) Immunofluorescence shows recombination (red) in Nell2 (green) OECs (arrow). F’-F’’’) Sox10Cre recombination occurs in sparse neurons in the main olfactory epithelium positive for HuC/D. G-G’’’) Sox10Cre recombination appears in sparse nonneuronal cells in the VNO that are negative for HuC/D.

Thus, based on the extensive Wnt1-Cre2 recombination in non-neural crest-derived tissues, which has also been recently reported by others (Gandhi, Du, Pangilinan, & Harland, 2024), we decided to explore the *Sox10*-Cre transgenic mouse line as an alternative strategy to conditionally ablate *Yap*/*Taz* in the cranial NCCs (Jacques-Fricke, Roffers-Agarwal, & Gammill, 2012).

We validated the *Sox10*-Cre mice with Ai14 Rosa26 reporters and collected *Sox10*-Cre^Tg^/Ai14 embryos at E14.5 and postnatal stages (Supplementary Figure 1). Observation of nose sections showed expected recombination in the NCC-derived frontonasal cartilage and nasal mesenchyme (Fig. 4D). The OBs and brain showed no recombination except for presumptive pericytes (Fig. 4D,E”’ and Supplementary Fig. 1). Immunostaining against the OEC marker NELL2 confirmed the recombination in the OECs along the olfactory axons proximal to the OB (Fig. 4E’-E”’).

However, in line with previous studies using other presumptive NCC Cre tracing lines such as *Wnt1*-Cre, *Pax7*-Cre, and *P0*-Cre, we also found recombination in sparse cells in the OE and VNO (Fig. 4D,E”’ and 4F’-G”’). Some traced cells were positive for the neuronal markers HuC/D, while others were negative. At E14.5 we estimate that *Sox10*-Cre recombination accounts for less than 1% of the cells in the OE and VNO. In postnatal animals, less than 1% of the cells in OE and VNO were positive for tracing (Supplementary Fig. 1). Notably in the OE of postnatal animals, we identified traced sparse neuronal cells, and presumptive cells of Bowman’s glands.

### *Yap*^cHet^;*Taz*^cKO^ mutants have a reduced number of *Sox10*-positive cells along the developing olfactory fibers and a reduced number of Olfactory Marker Protein (OMP) positive neurons in the olfactory epithelium. (fig5)

Past investigations have conditionally knocked out *Yap* and *Taz* in presumptive NCCs using both the first-generation *Wnt1*-Cre (Danielian, Muccino, Rowitch, Michael, & McMahon, 1998) and the second-generation *Wnt1*-Cre^Sor^/*Wnt1*-Cre2 (Lewis, Vasudevan, O’Neill, Soriano, & Bush, 2013) as pre-migratory NCC genetic entry points (J. Wang et al., 2016). Homozygous loss of *Yap* and *Taz* in NC derived cells caused embryonic lethality at E10.5, making these genetic models unsuitable for our study (J. Wang et al., 2016). As we observed extensive recombination in the Wnt1-Cre2 lines we generated and analyzed *Sox10*-Cre/Yap^Flx/WT^;Taz^Flx/Flx^ (*Yap*^cHet^;*Taz*^cKO^).

Analyzing Sox10-Cre lineage tracing in postnatal animals indicated recombination in the Bowman glans and sparse neurons, which we estimated to be less than 1% (Supplementary Fig. 1). Gross observations at E14.5, in line with what previously reported for *Wnt1*-Cre/Yap^Flx/WT^;Taz^Flx/Flx^; revealed no noticeable morphological differences between control and mutant embryos (J. Wang et al., 2016). To understand whether the SCP derivatives in the nasal area are affected by altered *Yap* and *Taz expression*, we performed immunostainings against Sox10 on parasagittal sections of the head of both triple mutant and control WT embryos at E14.5. Quantifications of Sox10-positive cells proximal to the *lamina propria* of the olfactory epithelia and along the peripherin positive fibers of the olfactory sensory neurons (OSNs) revealed an overall ∼30% reduction in the number of Sox10-positive cells in the frontonasal region of *Yap*^cHet^;*Taz*^cKO^. However, performing more targeted quantifications revealed notable differences in the number of OECs in different head regions. Specifically, in the *Yap*^cHet^;*Taz*^cKO^ mutants, we found that, at this stage, the most dramatic reduction in Sox10-positive OECs was around the olfactory bulbs (-44%), which starkly contrasted with the much lower (-19%) reduction in Sox10-positive OECs lining the lamina propria of the olfactory epithelia (Fig. 5-D).

**Figure 5.**
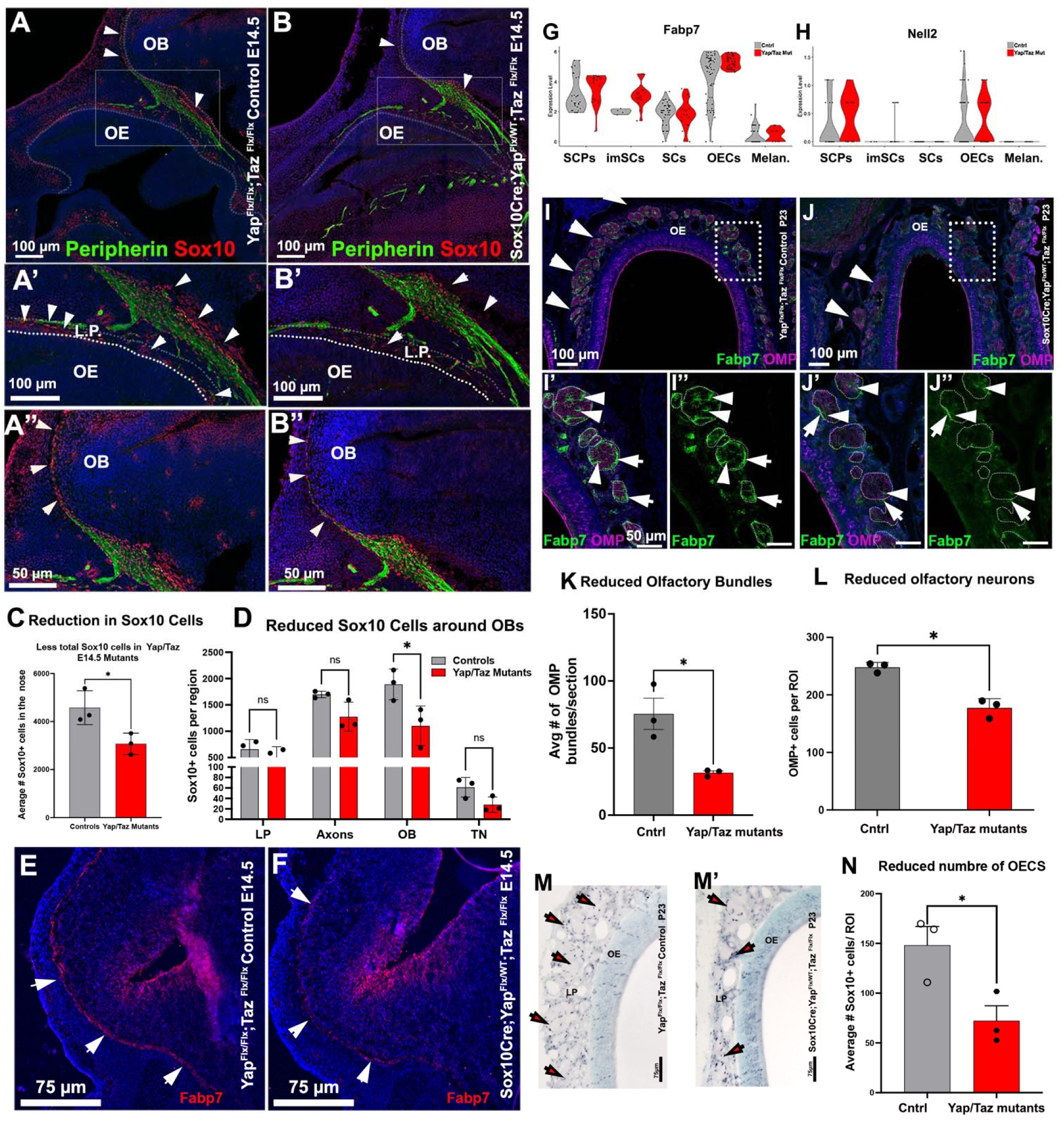
SoxCre/ Yap^Flx/WT^;Taz^Flx/Flx^ have reduced nasal glial cells and less olfactory neurons. A-B”) Immunofluorescence against Sox10 (red) and Peripherin (green) shows the nasal glia (arrowheads) associated with the olfactory-vomeronasal axons projecting to the Olfactory bulb; controls (A-A’) and SoxCre/Yap^Flx/WT^;Taz^Flx/Flx^ (B-B”). C) quantification showing a reduction in the total number of Sox10+ cells in conditional cKO (+/- SEM). D) Quantification indicates a statistically significant reduction of the olfactory ensheathing cells (OECs) around the olfactory bulbs (OBs) (+/- SEM). E-F) Immunoreactivity against Fabp7 of the OECs around the OB in controls (E) and mutants (F) where Fabp7 was barely detectable. G-H) Violin plots show comparable mRNA expression for Fabp7 and Nell2 in the OECs of control and mutant mice. I-J’) Fabp7 (green) and OMP (magenta) staining show fewer olfactory bundles around the OE (arrows in I and J; quantifications in (K) and reduced reactivity for both Fabp7 in the OECS surrounding the OMP+ axonal bundles in the lamina propria (I’-J”) (+/- SEM). L) OMP+ cell quantifications suggest a reduction in the olfactory neurons in SoxCre;Yap^Flx/WT^;Taz^Flx/Flx^ mutants (+/- SEM). M,M’) Sox10 immunostaining reveals a decreased number of OECs (black dots) in the lamina propria, and (N) quantification indicates a reduction in the number of olfactory neurons in SoxCre/Yap^Flx/WT^;Taz^x^mutants (+/- SEM).

Fabp7 immunostaining showed that Fabp7 was barely detectable in the *Yap*^cHet^;*Taz*^cKO^ mutants both at embryonic and postnatal stages (Fig. 5E-F), while it was evident in the control littermates. However, when comparing single-cell sequencing data from control littermates and *Yap*^cHet^;*Taz*^cKO^, we surprisingly found no significant changes in mRNA expression for *Fabp7* and *Nell2* mRNA in OECs (Fig. 5G,H). These findings suggest that the changes in Fabp7 immunodetectability either reflect variations in OECs number, translational or structural changes (Fig. 5C).

To determine whether further changes could be observed in postnatal animals, we analyzed P23 *Yap*^cHet^, *Taz*^cKO^ mice, and control littermates. Immunostaining for Sox10 revealed a persistent (∼50%) reduction in the number of OECs in *Yap*^cHet^;*Taz*^cKO^. In accordance with what was observed during embryonic development, BLBP immunostaining remained nearly undetectable in postnatal *Yap*^cHet^;*Taz*^cKO^. Quantification of the number of the olfactory bundles in the nasal area of P23 animals showed a significant reduction in bundle number. Quantifying OMP-positive cells (Fig. 5L) in the olfactory epithelium also revealed a significant reduction in the number of olfactory sensory neurons.

### *Yap* and *Taz* are crucial in controlling gene expression and developmental progression in the craniofacial neural crest. (Fig6)

To investigate how loss of *Yap* and *Taz* altered gene expression during the development of migrating neural crest cells, we performed single-cell RNA sequencing of dissociated noses isolated from *Yap*^cHet^;*Taz*^cKO^ and control E14.5 embryos. When plotted, the cells from *Yap*^cHet^;*Taz*^cKO^ and control groups formed grossly comparable Uniform Manifold Approximation and Projection (UMAP) clusters across genotypes, as shown in Figures 6A-A’.

**Figure 6.**
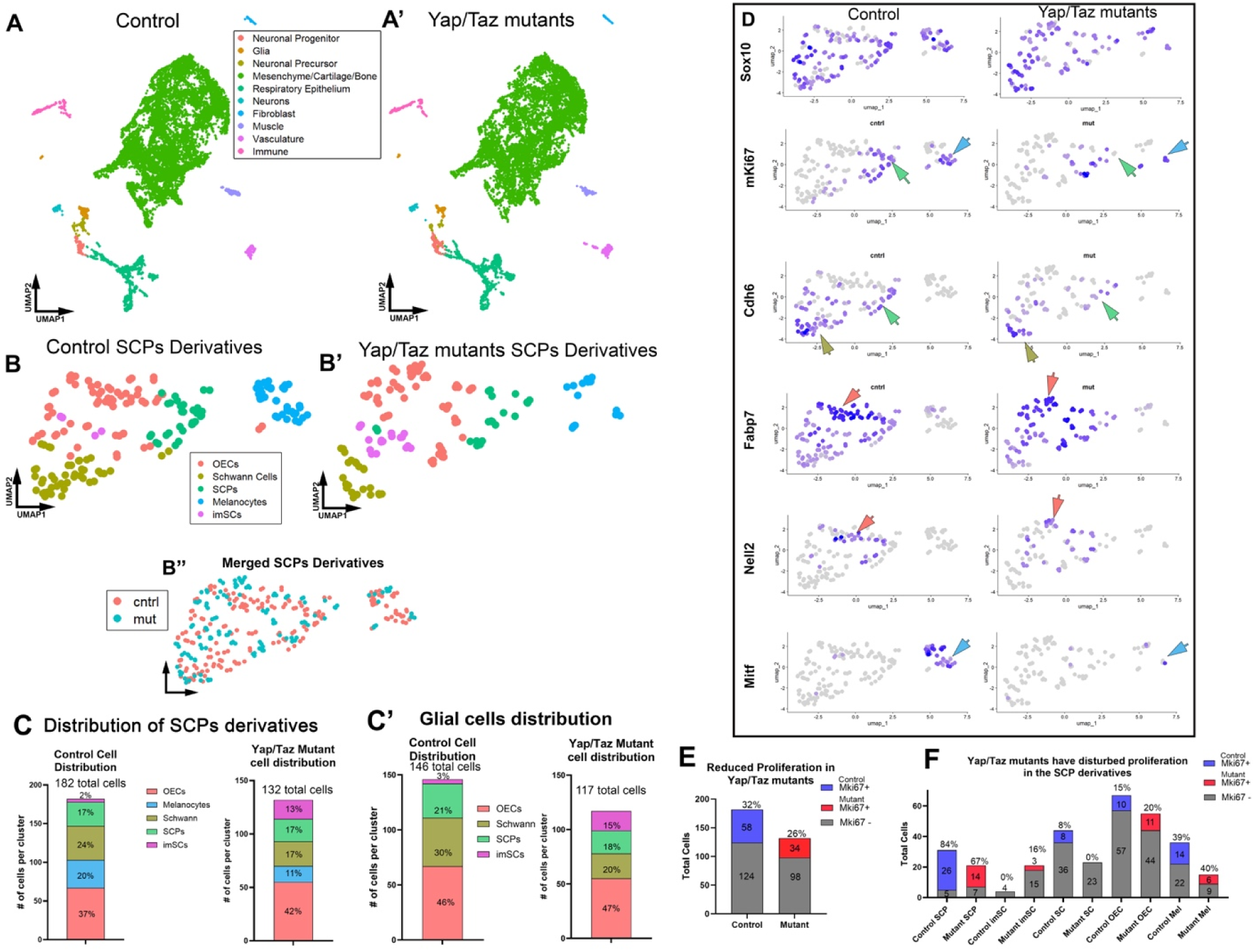
Sc-Seq of control and YAP/TAZ mutants shows defective development of the SCP derivatives. A-A’) UMAPs of control and mutant form comparable cell clusters (cell types indicated in the legend). B,B’) Subcluster of Sox10+ presumptive SCP derivatives in controls. B’) The overlay of the UMAP of the control (orange) and mutant (tile) shows that mutant cells are distributed in the whole cluster. C) Different distribution of Sox10 cellular types (including melanocytes); the mutants gave a smaller yield of Sox10+ cells and had a larger representation of immature SCs and OECs and a decrease in the percentage of SCs and melanocytes compared to controls. C’) The YAP/TAZ mutants have fewer Sox10+ glial cells; however, within the population, there is an increased representation of immature SCs and reductions in SCs, while the relative representation of OECs is similar to controls. D) Feature plots showing the expression of indicated genes. mKi67 shows a reduction in proliferative SCPs (green arrow) and melanocytes (light blue arrow), and Cdh6 shows a reduced number of presumptive SCPs (light green) and SCs (olive). E) YAP/TAZ mutants have reduced proliferative Sox10+ cells. The distribution of proliferative cells across the SCP derivative populations indicates a reduction in proliferation in the mutant SCPs, SCs, melanocytes and SCs but an increase in proliferating imSCs.

For both genotypes, we identified presumptive SCPs and their presumptive *Sox10*-positive derivatives: SCs, OECs, and melanocytes (Fig. 6B-B’; D). Aligning with our observations from counting the number of SOX10-positive cells, we found a consistent ∼27% reduction in the total number of cells expressing *Sox10* mRNA (Fig. 6C-C’). Among the *Sox10*-expressing cells, the percentage of SCPs remained unchanged (Fig. 6 C-C’), but we found a dramatic increase in the representation of immature SCs (from 2% in controls to 13-15% in the *Yap*^cHet^;*Taz*^cKO^ mutants), which is suggestive of changes in SC maturation dynamics. Focusing on the SCPs derivatives (Fig. 6C), we noted a ∼50% reduction in putative melanocytes and a ∼30% reduction in cells expressing SC identity markers. Surprisingly, though reduced in number, we found no gross changes in the relative percentage of cells with putative OEC identity.

By analyzing the number of Sox10 proliferative cells (*Mki67*+), we found a reduction in the number of cells undergoing mitosis in the Yap^cHet^; Taz^cKO^ mutants (Fig. 6E,F). Specifically, by examining the distribution of proliferative cells across different cell types (Fig. 6F), we found evidence of a dramatic reduction in the percentage of proliferating SCPs (-17%). In line with other studies, we observed a complete loss of proliferative SCs (Belin, Zuloaga, & Poitelon, 2017; Jeanette et al., 2021). Notably, we found an increase in the number and proportion of proliferative immature Schwann Cells, suggestive of a delayed transition from imSCs to SCs. As reported by others, we also found a dramatic loss of melanocytes (Manderfield et al., 2014), while the less affected cells, though reduced in number, appeared to be the proliferative OECs.

To gain a better understanding of the impact of the identified gene changes (Fig. 7) on key cell pathways, we performed a predictive analysis of pathway activity and regulatory networks using Qiagen’s Ingenuity Pathway Analysis tool (Fig. 8). Analysis of the data indicates that the SCPs show an increase in the expression of genes involved in DNA Methylation and Transcriptional Repression, an increase in Rho GTPase activity, changes in TGF-b and Notch signaling, a reduction in G2-M phase during mitotic metaphase, and an increase in ubiquitination (Fig. 7A, 8A). The immature SCs are amongt the cells more affected in terms of cell distribution (Fig. 6C,C’) and were found to be the cell state with the largest group of downregulated genes (Fig 7B;8B). These included genes involved in DNA synthesis, mRNA processing, decrease in ubiquitination, and Golgi to ER trafficking, suggestive of defective protein synthesis. In the SCs (Fig 7C,8C), we observed a decrease in RUNX2 activity, DNA synthesis, and Wnt1 signaling, while in the OECs, we noted a decrease in RNA Pol II transcription and genes involved in mRNA processing, which suggests that several defects might arise from changes in translation. In line with previous studies, we found a loss of *Mitf* (Manderfield et al., 2014) and *Mitf*-dependent gene expression in the melanocytes including cKit (Fig. 7E-8E). In supplementary (Supl.) Tables 1-5 provide the complete list of differential gene expression (DGE) between control and mutant samples, while the genes affecting different pathways are detailed in Supl. Tables 6-10.

**Figure 7.**
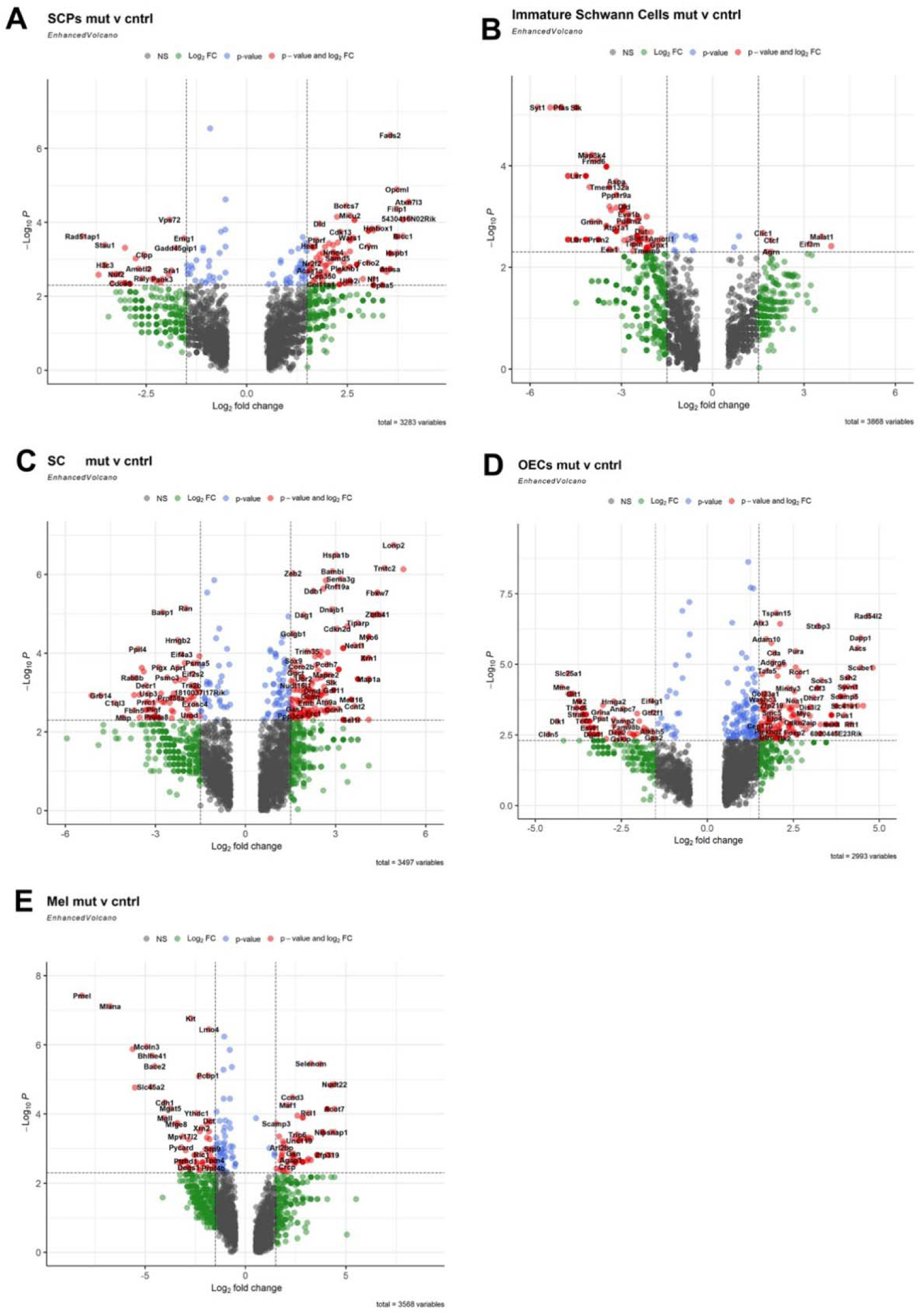
A-E) Volcano plots displaying differentially expressed genes, between controls (left) and conditional Yap-Taz mutants (right), in Schwann cell precursors (A), immature Schwann cells (B), Schwann cells (C), olfactory ensheathing cells (D), and melanocytes (E). Red dots indicate p-value ≤ 0.05 and fold change ≥ 1

**Figure 8.**
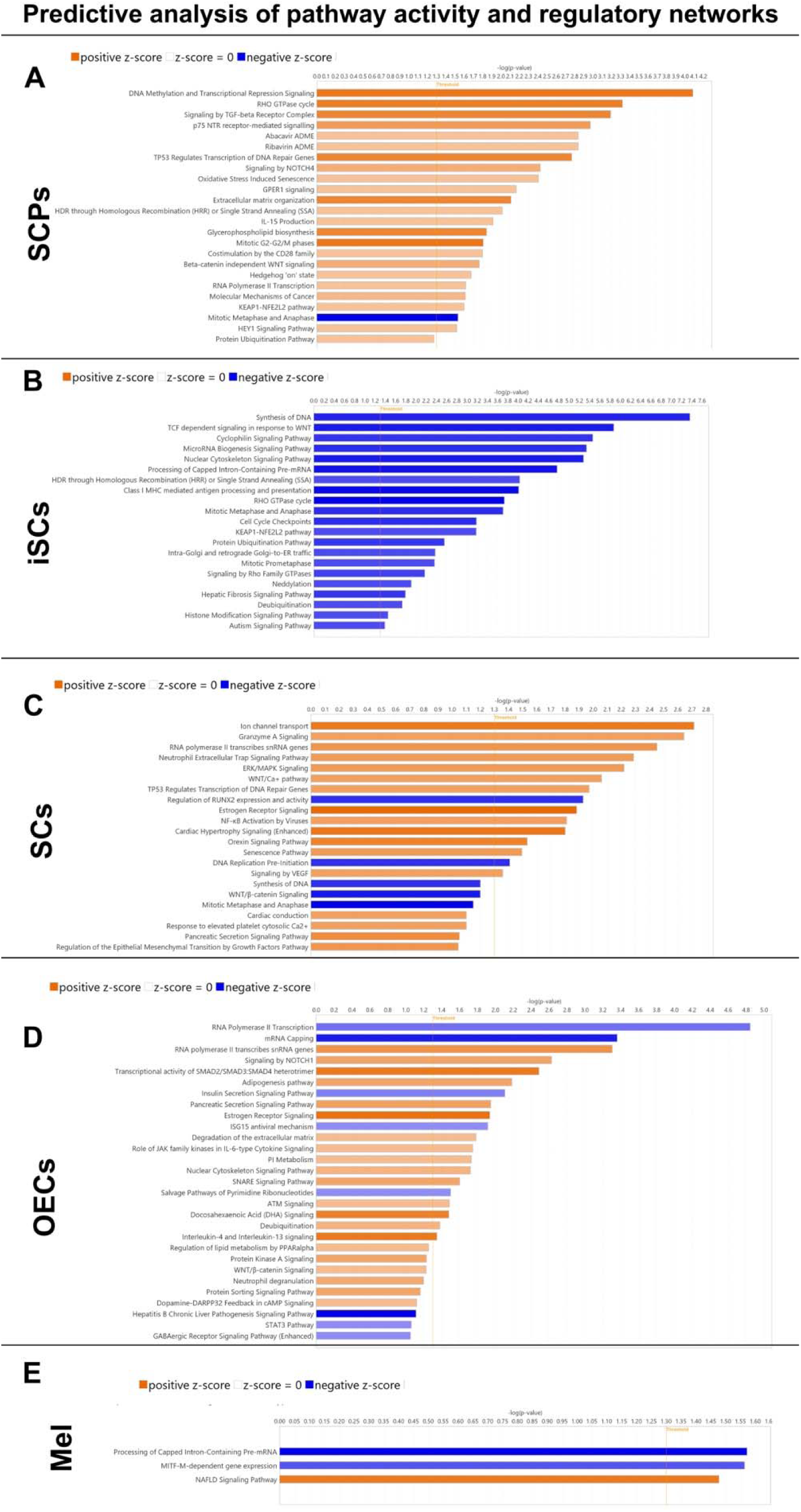
A-E) predictive analysis of changes in pathway activity and regulatory networks using Ingenuity Pathway Analysis(QIAGEN) in Schwann cell precursors (A), immature Schwann cells (B), Schwann cells (C), olfactory ensheathing cells (D), and melanocytes (E) of Yap Taz mutants and control from the differential gene expression data performed in Figure 7.

### *Yap*^cHet^;*Taz*^cKO^ mutants have a comparable distribution of GnRH-1 neurons to the control group (Fig.9)

Previous studies have shown that genetic mutations compromising neural crest development and the development of OECs can disrupt the migration of GnRH-1 neurons (Barraud et al., 2013; Pingault et al., 2013). After observing a reduction in OECs in the nasal region of *Yap*^cHet^;*Taz*^cKO^ mutants, we felt decided to investigate potential secondary effects on GnRH-1 neuronal migration. Contrary to our expectations of disrupted migration, anti-GnRH-1 immunostaining and quantifications of GnRH-1 neuron numbers and distribution showed no obvious differences between genotypes at the observed developmental stage (Fig. 9).

**Figure 9.**
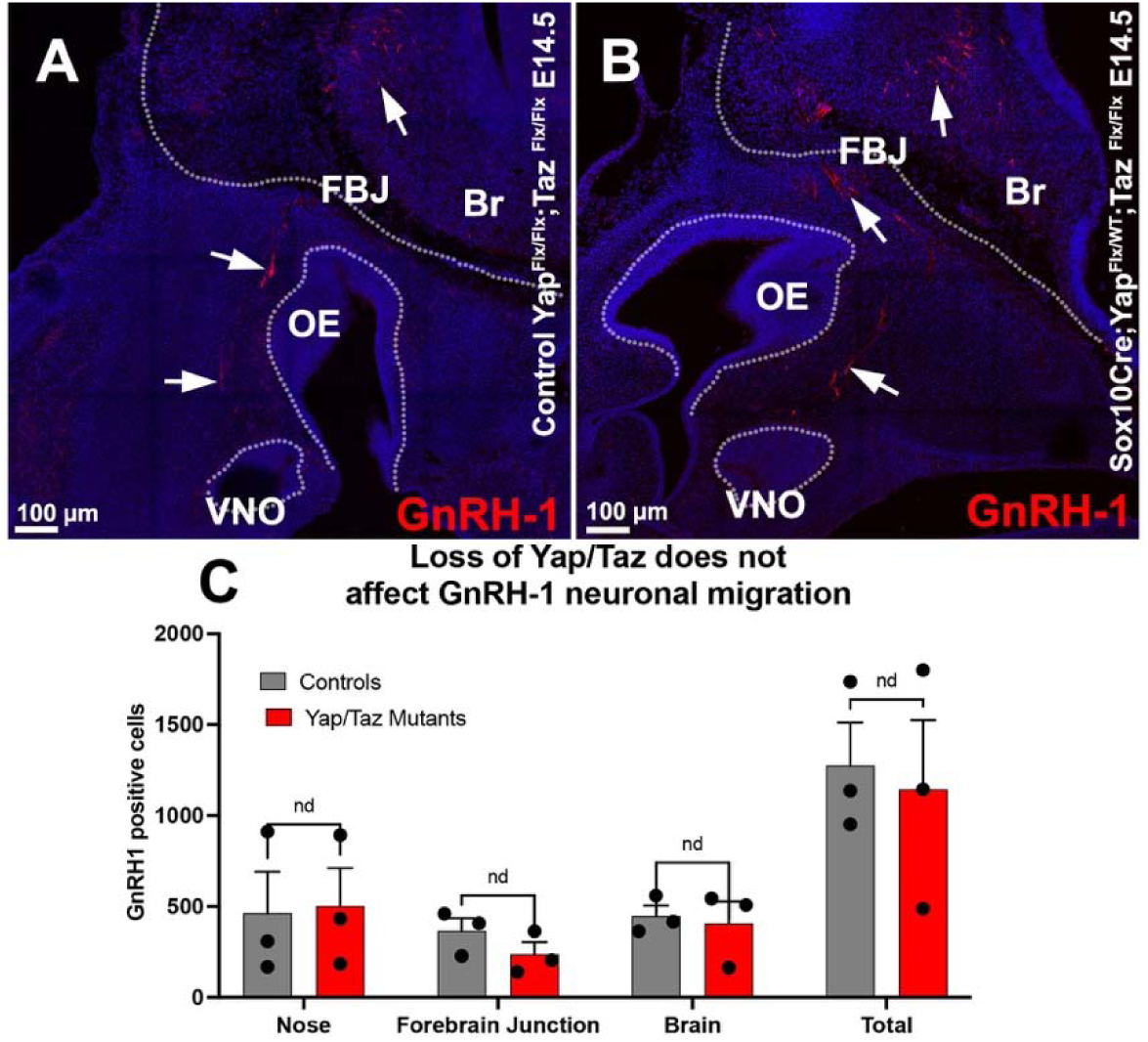
A) Immunofluorescence anti-GnRH-1 shows GnRH-1 neurons migrating to the brains of control and YAP/TAZ mutants. B) Quantification of GnRH distribution between the nasal area, forebrain junction, and brain shows no significant difference in migrattion dynamics or total cell number across genotypes (+/- SEM; n=3).

### *Yap*^cHet^;*Taz*^cKO^ have a reduced number of vomeronasal neurons, a reduced number of OECs around the VNO, and defective vascularization of the VNO

The development and maturation of the olfactory epithelia are profoundly controlled by inductive signals from the surrounding nasal tissues, frontonasal mesenchyme, and OECs, which are of NC origin (Forni et al., 2013; LaMantia, Bhasin, Rhodes, & Heemskerk, 2000; Perera et al., 2020). Based on this we analyzed the vomeronasal neurons in *Yap*^cHet^;*Taz*^cKO^ mice. The VNO mainly comprises two types of neurons: apical, which expresses the transcription factor Meis2 and the V1R family of receptors, and basal vomeronasal sensory neurons (VSNs), which express the transcription factor Tfap2e (AP-2e) as they mature and the V2R family of receptors (Katreddi & Forni, 2021) (Fig 10A-B’). Sox10Cre lineage tracing suggests the existence of Sox10 expressing progenitors which give rise to only sparse neurons (less than 1%) in the vomeronasal area (Supplementary Fig. 1). However, quantifications showed, as for the main olfactory epithelium a significant reduction in vomeronasal neurons (Fig. 10A-C).

**Figure 10.**
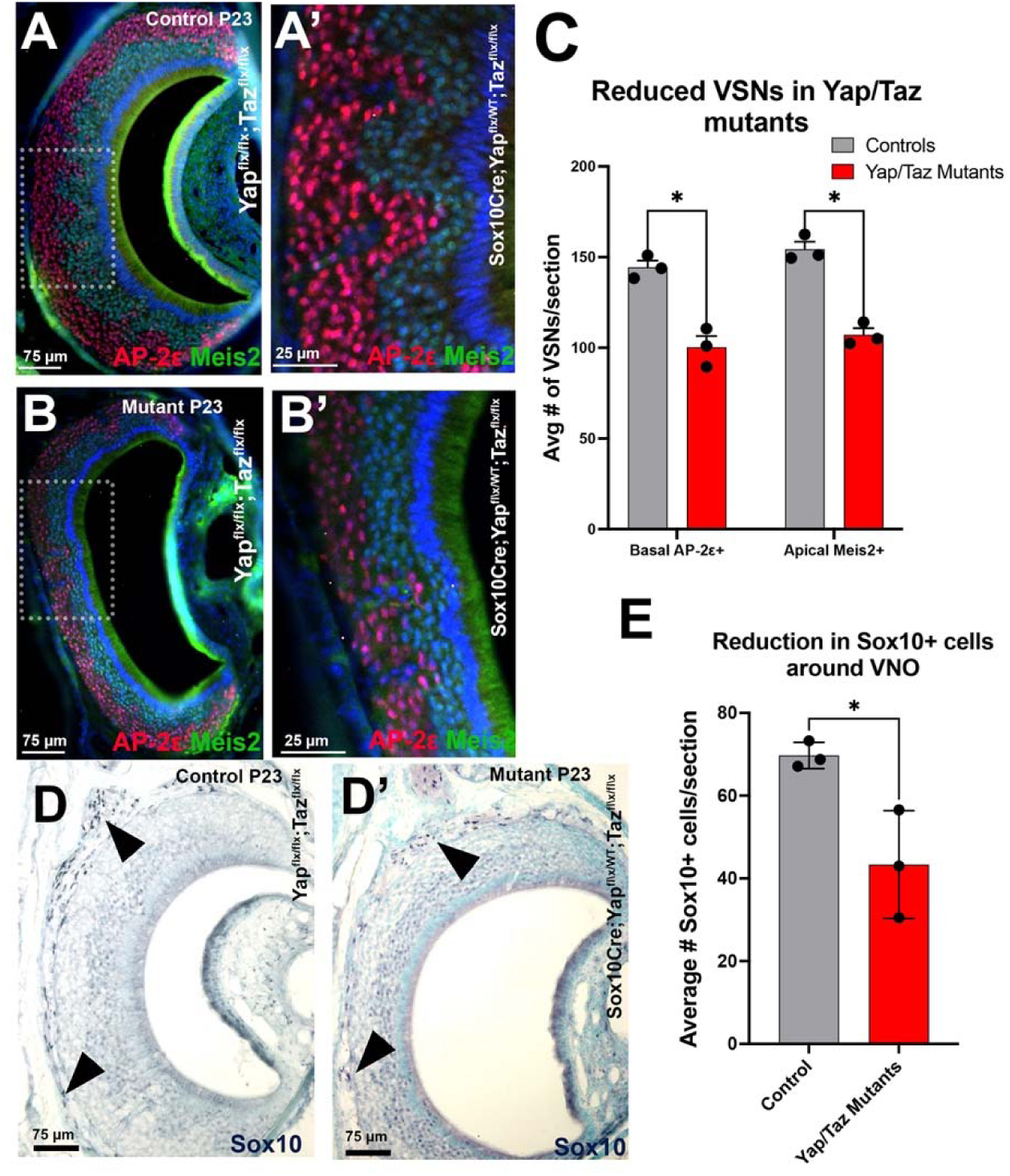
Postnatal Sox10 Cre/Yap^Flx/WT^; Taz^Flx/Flx^ have a reduced number of vomeronasal sensory neurons, reduced number of OECs and vasculature. P23, immunostaining against AP-2e (red) and Meis2 (green) in Controls (A-A’) and Yap Taz mutants (B-B’) shows a decreased thickness of the epithelium and visibly fewer apical Meis2+ and basal AP-2e neurons. A’ and B’ are enlargements of the boxed areas in A, B. C) Quantifications show a reduced number of apical and basal VSNs in the mutants (red) (+/- SEM). D, D’ Sox10 immunostaining in nickel DAB reveals Sox10+ cells in the lamina propria of the VNO (arrows) of controls D and Yap/Taz mutants D’. E) Quantifications show a significantly reduced number of Sox10+ cells in the conditional Yap;Taz mutants (+/- SD).

These data indicate defective development of NCC-derivative in the nasal area. The nasal mesenchyme and OECs have broad secondary effects on the number of chemosensory neurons, likely resulting from changes in local inductive signals. Quantification of the OECs along the basal lamina and within the lamina propria of the VNO indicated a reduced number of glial cells (Fig. 10D-E).

### The reduction in basal/V2R vomeronasal neurons decreases the number of vascular intrusions (Fig.11)

Conditional loss of *Yap* and *Taz* in NC derivatives was previously described as indirectly causing defects in blood vessel formation in developing embryonic heads (J. Wang et al., 2016). Notably, both OECs and SCs have been described to induce vascularization in their surrounding tissue environment (Taïb et al., 2022; X. Wang et al., 2022). To understand whether the Sox10Cre *Yap*^cHet^;*Taz*^cKO^ mice have vascular defects, we analyzed the highly vascularized vomeronasal chemosensory epithelium.

**Figure 11.**
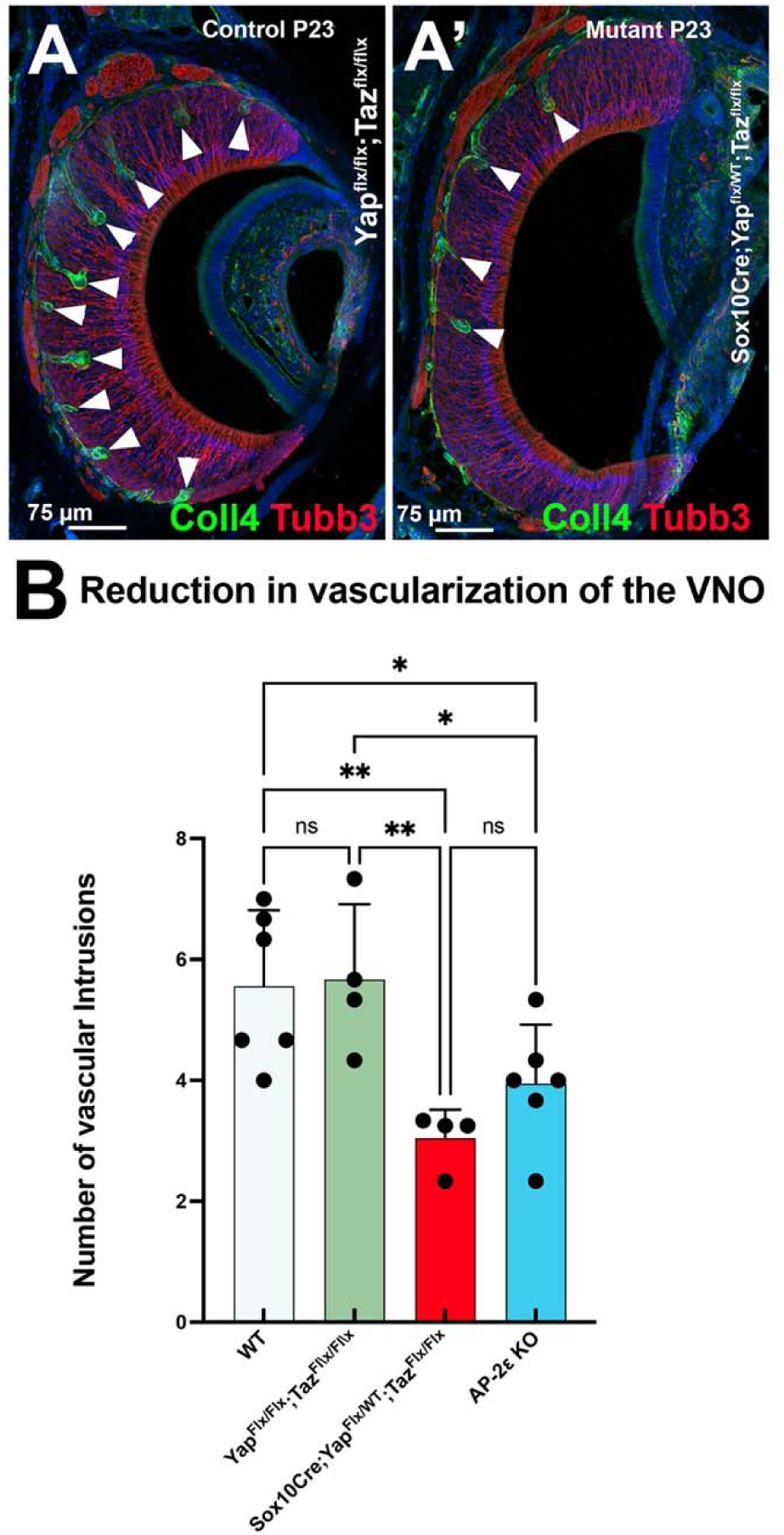
Reduction in basal VSNs correlated with a reduction in vascular intrusions. A-A’) Collagen IV (Col-IV) immunofluorescence highlights vasculature (arrows) in the VNO of controls Yap^Flx/Flx^; Taz^Flx/Flx^ and Sox10 Cre/Yap^Flx/WT^; Taz^Flx/Flx^ mutants at P23. B) Quantification of the number of vessels per VNO section demonstrates similar reduction in number of vascular intrusions in Yap Taz mutants and in AP-2eKOs

In the vomeronasal organ, blood vessels invade the basal territories where the basal VSNs, positive for Tfap2e/AP-2ε; V2R; Gao, reside (Naik et al., 2020). Collagen type IV (Col IV) is a basement membrane protein enriched during vessel morphogenesis (Gross et al., 2021). We previously described Col IV being expressed around the Pecam1+ vasculature, invading the basal regions of the vomeronasal organ (Naik et al., 2020). Quantifying the number of vascular intrusions in the vomeronasal organ of P23 mice revealed a significant reduction in the vascularization of the vomeronasal epithelium of mutant mice (Fig 11A,B).

The transcription factor Tfap2e is selectively expressed in the basal VSNs of the VNO, not in the vasculature nor the nasal mesenchyme (Hills et al., 2024; Lin et al., 2022; Lin et al., 2018). Tfap2e KO (AP-2e KO) mice have a reduced number of VSNs (Lin et al., 2018). To understand if such a phenotype could result from defects in the vomeronasal epithelium rather than from broad defects in neural crest-derived cells, we analyzed the number of vascular intrusions in AP-2e KO mice (Fig. 11B). Quantifying the number of vascular indentations in the basal territories of the VNO indicated similar vascularization reductions between Yap^cHet^;Taz^cKO^ and AP-2e KO mice. These data suggest that the vascular phenotype in Yap^cHet^;Taz^cKO^ mice is, at leat in part, secondary to the decrease in basal VSNs.

## Discussion

The nasal glial populations associated with the terminal nerve, olfactory, and vomeronasal neurons are commonly referred to as the OECs. Over the last thirty years, these cells have generated interest and excitement in the scientific community for their potential therapeutic uses after spinal cord lesions (Assinck, Duncan, Hilton, Plemel, & Tetzlaff, 2017; DeLucia et al., 2003; Goulart et al., 2016; Gu et al., 2017; Huang et al., 2024; Khankan et al., 2016; Ma et al., 2010; Raisman, 2001; Ramon-Cueto, Plant, Avila, & Bunge, 1998; Riddell, Enriquez-Denton, Toft, Fairless, & Barnett, 2004). However, the molecular characterization and developmental processes of these glial cells have only been partially addressed.

The OECs are essential for organizing the olfactory nerve bundles, establishing the olfactory system connections with the olfactory bulbs, and, crucially, ensuring the correct migration of GnRH-1 neurons along the fibers of the terminal nerve to the brain (Barraud et al., 2013; Pingault et al., 2013; Taroc, Naik, et al., 2020). Initial studies on the embryonic lineage of these cells proposed that they originate from the olfactory placode (Couly & Le Douarin, 1985). However, the neural crest origin of the OECs was then proposed and tested, over a decade ago, through a series of genetic lineage tracing in mice and quail-chicken tissue graft experiments (Barraud et al., 2010; Forni et al., 2011). Several genetic lineage tracing studies also proposed the contribution of the NCCs, or cells with genetic overlaps to the NC, to the olfactory epithelia (Barraud et al., 2010; Forni et al., 2011; Katoh et al., 2011; Koontz, Urrutia, & Bronner, 2022; Murdoch, DelConte, & Garcia-Castro, 2012; Saxena et al., 2013). These studies proposed that some OSNs, VSNs and microvillar cells can originate from NCCs. However, the reliability of tracing neural crest lineages using various genetic models has been a matter of debate (Aguillon et al., 2018; Barraud et al., 2013; Steventon, Mayor, & Streit, 2014).

Projections from the trigeminal nerve innervate the nasal region of rodents (Hummel & Frasnelli, 2019; Tremblay & Frasnelli, 2018; Ulusoy et al., 2022). Previous studies in rodents suggested the existence of heterogeneous populations of OECs in the nasal area of mice, with cells showing similarities to both SCs of the peripheral nervous system and other cells exhibiting specific OEC features (Guerout et al., 2010; Miedzybrodzki, Tabakow, Fortuna, Czapiga, & Jarmundowicz, 2006; Smith et al., 2020). However, to date, only a handful of studies have attempted to follow the development and identify the key molecular markers and mechanisms responsible for forming OECs (Perera et al., 2020; Rich et al., 2018). A recent study suggested that the origin of the OECs may differ from that of the SC, with the OECs forming from mesenchymal progenitors and SCs from SCPs. However, studies from the peripheral system suggest that glial and mesenchymal programs are divergent (M. E. Kastriti et al., 2022).

Our data, in line with a seminal study (Perera et al., 2020), show that both SCs and OECs are present in the nasal area. Moreover, we confirmed the previously described markers and identified new ones that allow us to discriminate between the developing olfactory ensheathing cells and SCs of the PNS (Fig. 2). By conducting single-cell sequencing of the developing nose, we identified and isolated Hub/SCPs cells (Stierli & Sommer, 2022) and preumptive SCs, OECs and melanocytes. RNA Velocity analysis suggests that the OECs, like the SCs originate form SCPs (figure 1,2).

The OECs appear as early as E11.5 when the olfactory migratory mass leaves the developing olfactory pit (Forni et al., 2011; Perera et al., 2020). If the SCPs giving rise to the OECs arrive at the developing olfactory system from other peripheral projections or they form from “hub cells” (Maria Eleni Kastriti et al., 2022) contributing to the nasal mesenchyme (Perera et al., 2020; Soldatov et al., 2019; Xie et al., 2019), is something that should be further investigated.

Differential gene expression analysis revealed specific marker genes for each of the cell population forming from Sox10 SCPs in the nose (Fig.2), making it possible to discriminate between populations based on differential protein expression. Further validation through immunofluorescent staining confirmed the specificity of these markers in different cell populations, making Nell2 and Fabp7 reliable OEC markers, while Cdh6 was found to be enriched in SCPs and SCs along the trigeminal projections.

The transcription factors Yap and Taz have previously been indicated to play a key role in modulating genes responsible for craniofacial and SCs development and function (Yannick Poitelon et al., 2016; J. Wang et al., 2016). Notably, we found that Yap and Taz, along with the co-factors of the Tead family, are also expressed in the nasal SCPs and their presumptive derivatives, including the OECs. Based on this, we decided to explore how much Yap-Taz play a similar role in OECs’ as described for the Schwann cells of the PNS.

Genetic tracing using the early neural crest pre-migratory marker Wnt1 suggested that neural crest cells may co-mingle with the cells of the developing olfactory placode and integrate within the olfactory placode. However, by testing the recombination of the widely used second generation of Wnt1-Cre mice (Wnt1-Cre2)(Lewis et al., 2013), we found broad recombination in tissues of non-neural crest origin, including the olfactory bulbs (Fig. 4). Notably, a recent study found recombination in endodermal cells (Gandhi et al., 2024). Wnt1 is not expressed in the cells of the developing nose (Fig. 4). However, in line with recent observations by others, we noted extensive recombination of Wnt1-Cre2 in non-NC derivatives, including most of the cells the olfactory epithelium and the olfactory bulbs. The extent of recombination renders this mouse line unsuitable for experiments aimed at selectively manipulating genes in the cranial neural crest. We, therefore, adopted Sox10 Cre as an alternative. This mouse line gave good recombination in the nasal mesenchyme and OECs (Fig. 4). However, consistent with tracing using the first generation of Wnt1-Cre mice (Forni et al., 2011; Katoh et al., 2011), we also found sparse traced cells in the olfactory and vomeronasal epithelia (Fig. 4), wihch estimated to be less than 1% of the olfactory/vomeronasal sensory neurons (Supplementary Fig. 1). Furthermore, consistent with zebrafish data (Saxena et al., 2013), we found neurons positive for Sox10 tracing in the main olfactory epithelium. Additionally, in line with previous observations (Perera et al., 2020), we confirmed Sox10 expression and recombination in Bowman’s gland progenitors. These data confirm the existence of some Sox10-expressing cells that contribute to the olfactory epithelia, if these are NC derivatives still remains an open question.

Using Sox10Cre as a genetic entry point, we generated Sox10Cre *Yap*^cHet^;*Taz*^cKO^. The overall number of OECS and other Sox10 cells associated with neurons in the nasal area of Yap^cHets^/Taz^cKOs^ was significantly reduced both in embryonic and postnatal noses (Fig. 5,6,10). Manipulation in Yap and Taz affected the proliferation of the SCPs, leading to reduced formation of their derivatives and likely altered maturation, as suggested by the accumulation of cells in an imSCs state (Fig. 6). Notably, defective proliferation was also noted in SCs and melanocytes. Among the SCPs derivatives forming in the nose, Sc-seq showed extensive decreases in gene expression and an accumulation of cells in the immature SCs stage (Fig. 6,7,8; Supl. Tables 1,6). In the conditional mutant, we found a dramatically decreased percentage of cells progressing into the Schwann cell and melanocyte differentiation program (Fig. 6).

Pax3, together with Sox10, controls the expression of the microphthalmia-associated transcription factor (Mitf), which is essential for melanogenesis. Mutations in Sox10 and Pax3 in humans lead to Waardenburg syndrome, marked by pigmentation abnormalities and hearing loss resulting in OEC defects. Taz/Yap65 interacts with Pax3 to regulate Mitf expression(Manderfield et al., 2014). Manipulation of migratory/post-migratory Taz/Yap using Sox10-Cre confirmed the loss of c-KIT and Mitf (Manderfield et al., 2014) and downstream melanocyte genes (Fig 7E-E3; Supl. Table 2,7), as previously reported after pre-migratory *Wnt1*-Cre recombination (Manderfield et al., 2014). Along with that we found differential changes in gene expression and signaling pathways in all analyzed maturation stages of the examined SCP derivatives (Fig. 7,8; Supl. Tables 5,10).

Development of the craniofacial mesenchyme, cartilage, bones and the the olfactory epithelial are intimately interdependent. In fact, the differentiation in the olfactory epithelium depends on inductive interactions with the surrounding neural crest-derived tissues (Forni et al., 2013; LaMantia et al., 2000). Our histological analysis of *Yap*^cHet^;*Taz*^cKO^ found significant decrease in the number of olfactory and vomeronasal neurons, which are mostly negative for Sox10-Cre mediated recombination (Fig. 5, 10). Notably, in a previous study using Ascl1 KO, we observed that changes in the number of the OECs are directly dependent on the number of olfactory neurons (Taroc, Naik, et al., 2020). In *Yap*^cHet^;*Taz*^cKO^ we found that the neurons of the VNO are reduced by ∼30% (Fig. 10). So, due to the extent of NCC’s contribution to the nasal area, the role of the nasal mesenchyme in controlling the olfactory structure’s development, and the role of olfactory/vomeronasal neurons in controlling the OECs, it is virtually impossible, with the experimental paradigm we adopted, to discern between cell-autonomous and secondary changes on the OECs development.

Previous studies have also shown that the development of the OECs is crucial for olfactory development and GnRH-1 neuronal migration (Barraud et al., 2013; Pingault et al., 2013; Zhou et al., 2017). Our observations on Yap and Taz mutants showed that, although significantly reduced in number, the defective OECs development did not affect the GnRH-1 neuronal migration to the brain despite the loss of function in Yap and Taz (Fig. 9). Notably, evidence indicates that, despite the syndromic manifestation of olfactory and GnRH migratory defects in Kallmann syndrome, the development of the olfactory system and the terminal nerve-GnRH system overlap only partially (Amato et al., 2024; Taroc, Prasad, Lin, & Forni, 2017). Our data, consistent with observations in Frzb mutants (Rich et al., 2018), suggest that the TN-GnRH-1 system is highly resilient in its ability to access and invade the brain. Additionally, the role of the OECs in regulating olfactory and TN development may depend on distinct mechanisms, if not on different cellular processes altogether.

The basal territory of the vomeronasal organ of mice, where the V2R-expressing neurons reside, is highly vascularized with blood vessels forming regular indentations in the sensory epithelium (Dietz, Senf, & Neuhaus, 2025; Katreddi & Forni, 2021; Naik et al., 2020). The *Yap*^cHet^;*Taz*^cKO^ mutants have significantly fewer (around -30%) vomeronasal organ neurons (Fig. 10) and fewer vascular indentations than the controls (Fig. 11). To understand whether the changes in vascularization in the VNO were secondary to defects of the vomeronasal organ, we analyzed the vascularization of the VNO of Tfap2e/AP-2e KO mouse mutants(Lin et al., 2022; Lin et al., 2018). These mice have dramatic defects in basal/V2R vomeronasal neurons but normal V1R neurons. AP-2e KO mice showed a comparable reduction in number of vascular indentations as the *Yap*^cHet^;*Taz*^cKO^. As AP-2e is not expressed in vasculature or nasal mesenchyme, these data suggest that loss of the basal/V2R neurons is sufficient to alter the number of vascular indentations of the basal territories of the VNO.

In conclusion, this paper provides compelling evidence indicating that OECs form from SCPs as the SCs of the PNS. Yap and Taz contribute to OECs development similarly to what was reported for the SCs. Moreover, we provide exciting evidence of the extent of the crosstalk between NC and placodal derivatives in controlling the development of the various cell types in the developing nose defining structures and homeostasis of the olfactory epithelia (LaMantia et al., 2000).

Our findings highlight the critical involvement of the Hippo Pathway in NC derivatives in shaping the nasal area, giving insights into potential underlying defects in the olfactory system associated with neurocristopathies and evolution (Jacobs, 2019).

## Material and Methods

### Animals

Sox10Cre (B6;CBA-Tg(Sox10-cre)1Wdr/J)(Matsuoka et al., 2005), Yap^Flx^, and Taz^Flx^ (*Wwtr1^tm1.2Hmc^* Yap^1tm1.2Hmc^/WranJ)(Reginensi et al., 2013); Wnt1Cre2 (B6.Cg-*E2f1^Tg(Wnt1-cre)2Sor^*/J) (Lewis et al., 2013); Rosa tdTomato; Ai14 (B6.Cg-*Gt(ROSA)26Sor^tm14(CAG-tdTomato)Hze^*/J) (Madisen et al., 2010)were purchased form JAX. Mouse lines were genotyped via PCR using the line-specific markers and indicated in Jax.org. Amplification products were analyzed by agarose gel electrophoresis. Animals were euthanized using CO2, followed by cervical dislocation. Mutant and wild-type mice of either sex were used. All mouse studies were approved by the University at Albany Institutional Animal Care and Use Committee (IACUC).

### Tissue preparation

Embryos were collected from time-mated dams where the emergence of the copulation plug was taken as E0.5. Collected embryos were immersion-fixed in 3.7% Formaldehyde/PBS at 4°C for 3-5 hours. Postnatal animals were perfused with ∼10mL of 3.7% Formaldehyde/PBS and then incubated overnight in 3.7% Formaldehyde/PBS at 4°C. All samples were then cryoprotected in 30% sucrose overnight, then frozen in O.C.T (Tissue-TeK) and stored at -80°C. Samples were cryosectioned (Embryos: Parasagittal; Postnatal heads: Coronal) using CM3050S Leica cryostat and collected on Superfrost plus slides (VWR) at 16 mm thickness.

### Immunohistochemistry

Primary antibodies and dilutions used in this study were: chicken-α-peripherin (1:1500, Abcam), rabbit-α-peripherin (1:2000, Millipore), SW rabbit-α-GnRH-1 (1:6000, Susan Wray, NIH), goat-α-olfactory marker protein (1:4000, WAKO), goat-α-AP-2ε (1:1000, Santa Cruz), mouse-α-Meis2 (1:500, Santa Cruz), mouse-α-Sox10 (1:1000, Sigma), goat-α-collagen IV (1:800, Millipore), rabbit-α-Fabp7(1:1000, Millipore), rabbit-α-Cdh6 (1:200, Invitrogen), sheep-α-Nell2 (1:250, R&D), mouse-α-Tuj1(1:1000, Abcam), mouse-α-HuCD-biotin(1:200, Molecular Probes), and goat-α-tdTomato(1:1000, Rockland). Antigen retrieval was performed in a citrate buffer prior to incubation with chicken-α-peripherin, goat-α-AP-2ε, mouse-α-Meis2, mouse-α-Tuj1, and mouse-α-Sox10. For immunoperoxidase staining procedures, slides were processed using standard protocols (Forni et al., 2013) and staining was visualized (Vectastain ABC Kit, Vector) using diaminobenzidine (DAB) in a glucose solution containing glucose oxidase to generate hydrogen peroxide; sections were counterstained with methyl green. For immunofluorescence, species-appropriate secondary antibodies were conjugated with Alexa-488, Alexa-594, or Alexa-680 (Molecular Probes and Jackson Laboratories) as specified in the legends. Sections were counterstained with 4′,6′-diamidino-2-phenylindole (1:3000; Sigma-Aldrich) and coverslips were mounted with Fluoro Gel (Electron Microscopy Services). Confocal microscopy pictures were taken on a Zeiss LSM 710 microscope and Zeiss LSM 980. Epifluorescence pictures were taken on a Leica DM4000 B LED fluorescence microscope equipped with a Leica DFC310 FX camera.

Images were further analyzed using FIJ/ImageJ software. Each staining was replicated on at least three different animals for each genotype.

### Cell quantifications

All embryonic image quantifications were performed on E14.5 parasaggital sections. Quantification of GnRH-1 neuronal distribution in mutant and control embryos was performed in the nasal region (VNO, axonal tracks surrounding the olfactory pits), forebrain junction, and brain (GnRH-1+ cells around the olfactory bulbs and were distributed within the forebrain). Number of cells were calculated for each animal as the average of the counted cells per series multiplied by the number of series cut per animal. Regional quantifications of Sox10 positive cells in control and mutant embryos was performed by counting positive Sox10 nuclei in the mesenchyme/basal lamina, associated with Peripherin positive axons before the forebrain junction, or within the forebrain junction/surrounding the olfactory bulbs. Total number of cells per region were averaged per animal. Quantification of Sox10 positive cells in post-natal animals were performed by counting the number of Sox10 postive nuclei in the surrounding lamina propria of both the main olfactory epithelium and the vomeronal epithelium. Total number of cells were averaged per animal. All significant differences were accessed by using unpaired t-Tests.

All post natal image quantifications were performed on P23 coronal sections. Post natal neuronal cell quantifications in the main olfactory epithelium and the vomeronasal organ were quantified using olfactory marker protein (OMP), and Tfap2e/Meis2 immunoflourescence respectively. Cells positive for the respective marker within the ROI (39,550 mm^2^) on each section were averaged per animal and per genotype. Vasculature quantification was performed by counting the number of Collagen IV positive intrusions with the vomeronal epithelium. Axon bundle quantifications were done on OMP stained sections. Number of bundles were averaged per animal and per genotype.

### Statistical analyses

All statistical analyses performed on image quantifications were done on GraphPad Prism 10 v10.4.0. Each quantification was performed on at least three biological replicates for control and mutant samples. Significant differences on all quantification were accessed using a Student’s unpaired t-test where significance was determined to be p value < 0.05.

### Single-cell RNA Sequencing

Single-cell RNA sequencing data used for Figures 1 – 5 was previously published by our lab (Amato et al., 2024). Newly generated single-cell RNA sequencing data (Fig. 6 – 8) was generated from the noses of E14.5 Sox10Cre/Yap^cHet^/Taz^cKO^ and control embryos. In short, the noses of embryos were dissected and isolated in cold 1x PBS. Single-cell dissociation was performed on each nose using dissociation solution (Papain in Neural Basal Media, Collegnase (0.5mg/mL), L-Cysteine (1.5 mM), and DNAse I (100u/mL)). Dissociated single-cells were stored in cell freezing media (90% DMSO in FBS) at -80C. Hindlimb or tail was collected from each embryo for DNA isolation for sex determination and genotyping through PCR. After determining the genotype and sex of each embryo, dissociated single cells were quickly defrosted in a 37C water bath. Cells were then washed and resuspended in 10% FBS in Neural Basal Media at a concentration of 700-1200 cells/mL. Automated cell counts were done on a T20 BioRad cell counter. GEM bead, cDNA, and library generation was performed using the 10X Chromium Next GEM Single Cell 3’ Reagent Kit v3.1 protocol with the 10X Genomics Chromium Controller located within the RNA Institute. Prepared libraries were then submitted to the Center of Functional Genomics for quality control assesment of the libraries and seqeuncing on the Illumina NextSeq2000 system.

FASTQ files were processed using 10X CELLRANGER v 9.0 and aligned to the mouse reference genome (GRCm39–2024-A) that is publicly available on the 10X website. The Cellranger output files were further processed using Velocyto to generate loom files. Velocyto loom files were then further analyzed using R/RStudio using the Seurat 5.1v package (Hao et al., 2024). To limit the inclusion of poor quality cells (eg. Dead cells, doublets) a filter for mito-genes expressed higher than 90%, and a filter for nFeature (genes) of more than 1000, and less than 6000 was used. Normalization of the dataset was performed using SCT Transform, and a Cell Cycle Scoring and Regression method available as a Seurat vignette was also performed to limit clustering based on cell cycle state. Seurat clusters were then converted to hda5 files in Seurat for use on Python. RNA velocity and pseudotime construction was done on Python using the package scVelo (Bergen, Lange, Peidli, Wolf, & Theis, 2020). Genes that were found to differentially expressed between mutant and control embryos were analyzed with the use of QIAGEN IPA (QIAGEN Inc., https://digitalinsights.qiagen.com/IPA) (Krämer, Green, Pollard, & Tugendreich, 2013).

## Data Availibility Statement

Single-cell RNA sequencing data sets used for analysis in this study have been uploaded and are available on the Gene Expression Omnibus (GEO) database, under GEO Accession number GSE286491. (https://www.ncbi.nlm.nih.gov/geo/query/acc.cgi?acc=GSE286491).

## Aknowledgments

This work was supported by the Eunice Kennedy Shriver National Institute of Child Health and Human Development (NICHD) under Grants 2R01HD097331-06A1 (P.E.F.) and 1R01HD114827-01A1 (P.E.F.), as well as by the National Institute on Deafness and Other Communication Disorders (NIDCD) under Grant R01HD097331 (P.E.F.). The Zeiss 980 microscope at the University at Albany was funded by the Office of the Director, NIH, under Award Number S10OD028600. Additional support was provided by the National Institute of Neurological Disorders and Stroke (NINDS) under Grants R01NS134493 (S.B.), R01NS110627 (Y.P.), and R03AG089077 (S.B. and Y.P.).

In situ hybridization images were sourced from the GenePaint database (https://gp3.mpg.de) (Visel, Thaller, & Eichele, 2004). Access and use of Qiagen’s Ingenuity Pathway Analysis software was provided by a floating license from NIH Library.

## Supplementary Figures/data

**Supplementary Figure 1.**
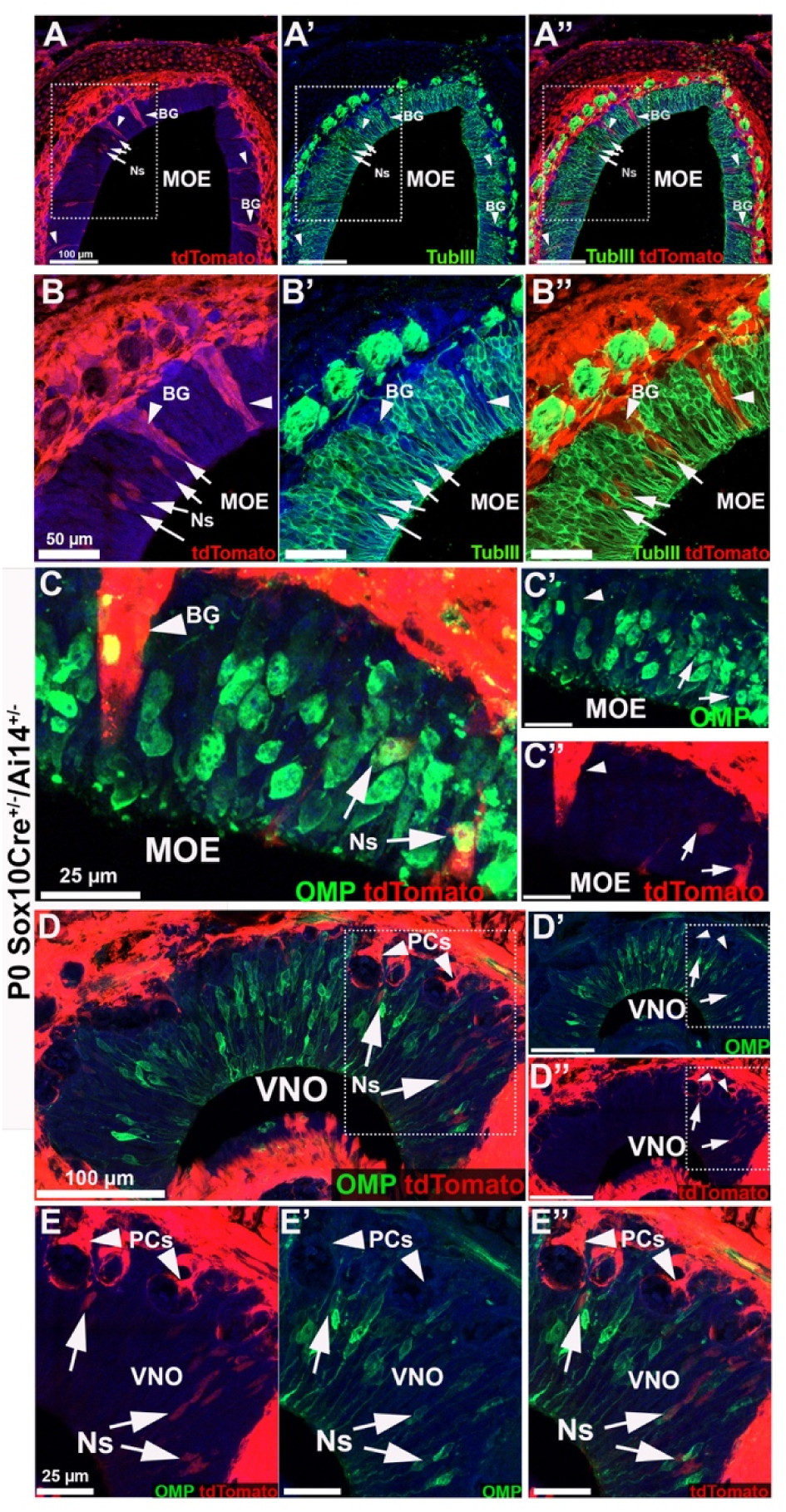
*Sox10*-Cre^Tg^/Ai14 mice tracing in postnatal mice reveals traced neurons in the central olfactory epithelium and vomeronasal organ. A-A’’, B-B’’’) P0 *Sox10*-Cre^Tg^/Ai14 immunofluorescence against neuronal marker Tub III (tubulin III, green), anti-tdTomato (red) and DAPI (blue) in the main olfactory epithelium (MOE). Recombination highlights neurons (Ns) positive for Tub III (arrows) and Bowman’s glands (BG). C-C’’) Immunofluorescent staining against the olfactory neuron marker OMP (olfactory marker protein, green) and anti-tdTomato (red) reveals traced neurons expressing OMP. D-D’’’, E-E’’’) Double immunofluorescent staining anti OMP and tdTomato within the VNO. Recombination can be observed in neurons positive for OMP and OMP negative pericytes (PCs) surrounding the vasculature.

## Legends for the supplementary tables provided as separate Excel files

**Supplementary Table 1: Differential Gene Expression of Immature Schwann Cell cluster between Yap^cHets/^Taz^cKOs^ and control.**

**Supplementary Table 2: Differential Gene Expression of Melanocytes cluster between Yap^cHets/^Taz^cKOs^ and control.**

**Supplementary Table 3: Differential Gene Expression of Olfactory Ensheathing Cell cluster between Yap^cHets/^Taz^cKOs^ and control.**

**Supplementary Table 4: Differential Gene Expression of Schwann Cell cluster between Yap^cHets/^Taz^cKOs^ and control.**

**Supplementary Table 5: Differential Gene Expression of Schwann Cell Precursor cluster between Yap^cHets/^Taz^cKOs^ and control.**

**Supplementary Table 6: Ingenuity Canonical Pathway Analysis of Immature Schwann Cell cluster between Yap^cHets/^Taz^cKOs^ and control.**

**Supplementary Table 7: Ingenuity Canonical Pathway Analysis of Melanocytes cluster between Yap^cHets/^Taz^cKOs^ and control.**

**Supplementary Table 8: Ingenuity Canonical Pathway Analysis of Olfactory Ensheathing Cell cluster between Yap^cHets/^Taz^cKOs^ and control.**

**Supplementary Table 9: Ingenuity Canonical Pathway Analysis of Schwann Cell cluster between Yap^cHets/^Taz^cKOs^ and control.**

**Supplementary Table 10: Ingenuity Canonical Pathway Analysis of Schwann Cell Precursor cluster between Yap^cHets/^Taz^cKOs^ and control.**

## Notes

### Competing Interest Statement

The authors have declared no competing interest.

